# Prior Information Shapes Perceptual Evidence Accumulation Dynamics Differentially in Psychosis

**DOI:** 10.1101/2025.07.03.662942

**Authors:** Léon Franzen, Sofia Eickhoff, Christina Andreou, Ioannis Delis, Julia Erb, Jens Kreitewolf, Rebekka Lencer, Claudia Lange, Lea-Maria Schmitt, Hannah Schewe, Niels Kloosterman, Sarah Tune, Stefan Borgwardt, Jonas Obleser

## Abstract

Humans rely on prior information to navigate sensory uncertainty: Such priors could shape the decision process before sensory evidence is gathered (origin model), or could amplify sensory evidence dynamically (gain model). Dysfunctionalities in the utilisation of priors may underlie hallucinatory percepts and delusional ideation in psychosis, yet their impact on decision-making across sensory modalities has remained unclear. Using a perceptual target-detection task across auditory and visual domains in laboratory and online samples, we applied hierarchical drift diffusion modelling to examine how prior probabilities shape evidence accumulation in clinical and non-clinical populations. We show that in healthy individuals, prior information enhances sensory gain and decision flexibility, as represented by drift criterion and rate, consistent with the gain model. In contrast, individuals with psychosis exhibit diminished sensory gain, relying instead on pre-evidence biases, consistent with the origin model. Notably, greater positive symptom severity predicted a reduction in traditional criterion decision bias. These results suggest that sensory gain deficits may serve as a computational marker for psychosis progression, linking dysfunctional prior use to perceptual aberrancies. By demonstrating how prior information modulates evidence accumulation across sensory modalities, our study advances the understanding of psychotic perception and decision-making, offering insights for computational psychiatry and fine-tuned clinical diagnostics.

## Introduction

Our daily environment is inherently noisy. The ability to interpret sensory information—such as “do I hear a human voice screaming or is it simply the wind howling?”—is critical for survival. To allow humans making decisions under uncertainty, incoming sensory evidence is integrated with prior knowledge ^1,2^. However, in psychosis, perceptual distortions and delusions emerge ^3^, suggesting fundamental disruptions in how prior knowledge interacts with incoming sensory evidence ^4–9^.

Generally, Bayesian theories of perception posit that observers combine prior knowledge with sensory evidence until a perceptual decision boundary is reached ^1,2,10–12^. In doing so, they exploit the variation in moment-to-moment available sensory evidence to optimise decision behaviour ^12–14^. However, the quality and context of sensory evidence matters, as prior expectations gain in weight when evidence is low, ambiguous or implicit ^11,15–17^. Theoretically, prior information can constrain incoming information in different ways: either before sensory evidence is being encoded and even presented (origin model ^13,18^), or by modulating encoding and evidence accumulation directly (gain model ^13,19,20^). Evidence exists for both mechanisms.

Speaking for an origin model, prior information has been shown to act before stimulus presentation by triggering existing stimulus templates in lower sensory cortex ^21^. This information elicits either downstream expectation-driven changes ^22^ or it biases the baseline activity of signal-selective units ^23^. Both mechanisms of the origin model are indicated by a starting point shift in the Drift Diffusion Model (DDM; ^13,18,24,25^).

In contrast, studies in line with the gain model report facilitatory effects of prior information during evidence accumulation. Under this account, prior information can facilitate how evidence is interpreted during evidence accumulation, reflected in changes of the DDM drift criterion (dc; ^26^) or in faster sensory processing speed, indicated by drift rate increases ^18,25,27–34^. Such modulations of evidence accumulation are arguably seen in changes in firing rate in parietal cortex of non-human primates when manipulating prior probabilities ^31^ and in changes in the evidence accumulation rate in humans ^28,34^.

Not least, simultaneous, dynamic changes in a priori bias (origin model) and in evidence accumulation (gain model) may be a third possible effect of probabilistic prior information, termed the multi-stage model ^35–38^. Such considerations highlight the multifaceted, not mutually exclusive, effects of prior information on visual perceptual decision-making. Though they do not answer conclusively whether probabilistic prior information rather affects an a priori bias or a facilitation of the evidence accumulation itself ^13,39^ within and across modalities.

When aiming for a generalisable answer, one must also consider decision modality, stimulus complexity, and trait-like characteristics of the person making these decisions. Investigations of this combination of factors remain scarce. Simple perceptual decision-making tasks point towards the evidence accumulation process (i.e., decision variable) exhibiting supramodal neural characteristics ^40–42^. However, contradictory behavioural and neural evidence casts doubts on purely supramodal characteristics ^19,43–47^. Behavioural data suggests that humans discriminate better in visual compared than auditory tasks ^43^, with reductions in signal-to-noise ratio (SNR) being most marked in the auditory modality ^46^. Also, two separate studies starting from similar assumptions, a recent manipulation of visual priors when watching degraded images ^48^ and an auditory study aiming for an analogous manipulation in degraded speech ^49^, speak against an unequivocally supramodal mechanism more reliant on prior knowledge. Thus, it remains elusive whether the evidence accumulation process itself exhibits strictly supramodal characteristics and whether supramodal characteristics extend to all decision components including a priori bias, sensory encoding, and the amount of required evidence.

While decision-making biases in psychosis have been studied, the computational mechanisms of how prior knowledge shapes perception across modalities remain poorly understood. Addressing this supramodal question requires within-subject studies using comparable stimuli in both auditory and visual domains—especially tasks that demand complex processing, akin to real-world conditions ^50^. However, most computational models of human perceptual decision-making have focused on a single-modality, typically simple visual tasks like random dot motion (e.g., ^41,45,51^) or on simultaneous multisensory inputs. Initial visual studies using sequential sampling models, which account well for perceptual decision-making, have reported changes in evidence accumulation speed in psychosis ^52,53^. Only recently has there been growing interest in applying drift diffusion models (DDMs) to auditory decisions ^54,55^. Further, these models have been scarcely applied in clinical populations and lack direct comparisons of audio-visual perceptual decisions within the same patients with psychotic disorders ^56^. Thus, cross-modal modeling studies with patients—especially those involving complex auditory tasks—are still rare but much needed. The current study aims to close this gap using a hierarchical DDM framework (HDDM; ^57,58^), which offers analytical benefits through group dependencies and accommodation of smaller samples.

Additionally, previous reports linking decision-making aspects to specific individual traits, which bear a relation to symptoms of psychosis, suggest that characteristics of the decision maker play a crucial role in disentangling the computational mechanisms of perceptual decisions. For instance, proneness to misperceptions in the form of hallucinations and schizotypal personality have been associated with aberrant weighting of sensory evidence in auditory decision-making ^59^. Similarly, increased schizotypy proneness has been reported to result in hyper-weighting of prior knowledge when given probabilistic cues ^60^. In contrast, delusion proneness has been associated with a reduced ability to update prior beliefs in response to changing contingencies in visual decision-making ^9,61^.

These traits are especially relevant for developing an encompassing understanding of the effects of prior information in aberrant perception. Individuals have difficulties overcoming existing biases ^62^ and those with psychosis frequently experience auditory and visual hallucinations, albeit with different prevalence ^63^, as well as delusions ^3^ Their hallucinations constitute false perceptual decisions and, following Bayesian and predictive coding accounts of psychosis ^6,7^ are believed to originate from a disbalance in the relative weighting of prior expectation and sensory evidence, particularly when the incoming sensory evidence is more ambiguous ^4,5,8,64–70^.

To investigate whether prior information acts supramodally in complex perceptual decision-making under high uncertainty, we combined auditory and visual tasks. We asked healthy participants and those with a diagnosis of psychosis to indicate whether they just heard a human voice in a sound or saw a human face in an image presented at one of two levels of noise, and to rate their confidence in the accuracy of their choice (Fig. 1A-D; see “Methods”). To manipulate prior information, we provided valid, probabilistic information on the amount of voice/face target stimuli in a given block of trials (Fig. 1B). This truthful prior information could either be informative (P^−^: fewer targets than half of the trials; P^+^: more targets than half of the trials) or uninformative (P^=^: half of the trials) for the binary detection decision (Fig. 1B). To separate perceptual aspects from strategic decision biases, our analyses focused on changes in complementary decision metrics including choice probabilities, as well as perceptual sensitivity (d’) and decision bias (criterion) from signal detection theory (SDT ^71^; Fig. 1E). Additionally, we exploited all available decision information by means of computational modelling of single-trial response times in mixed models and HDDMs (RTs; ^58,72^; Fig. 1F).

**Fig. 1.**
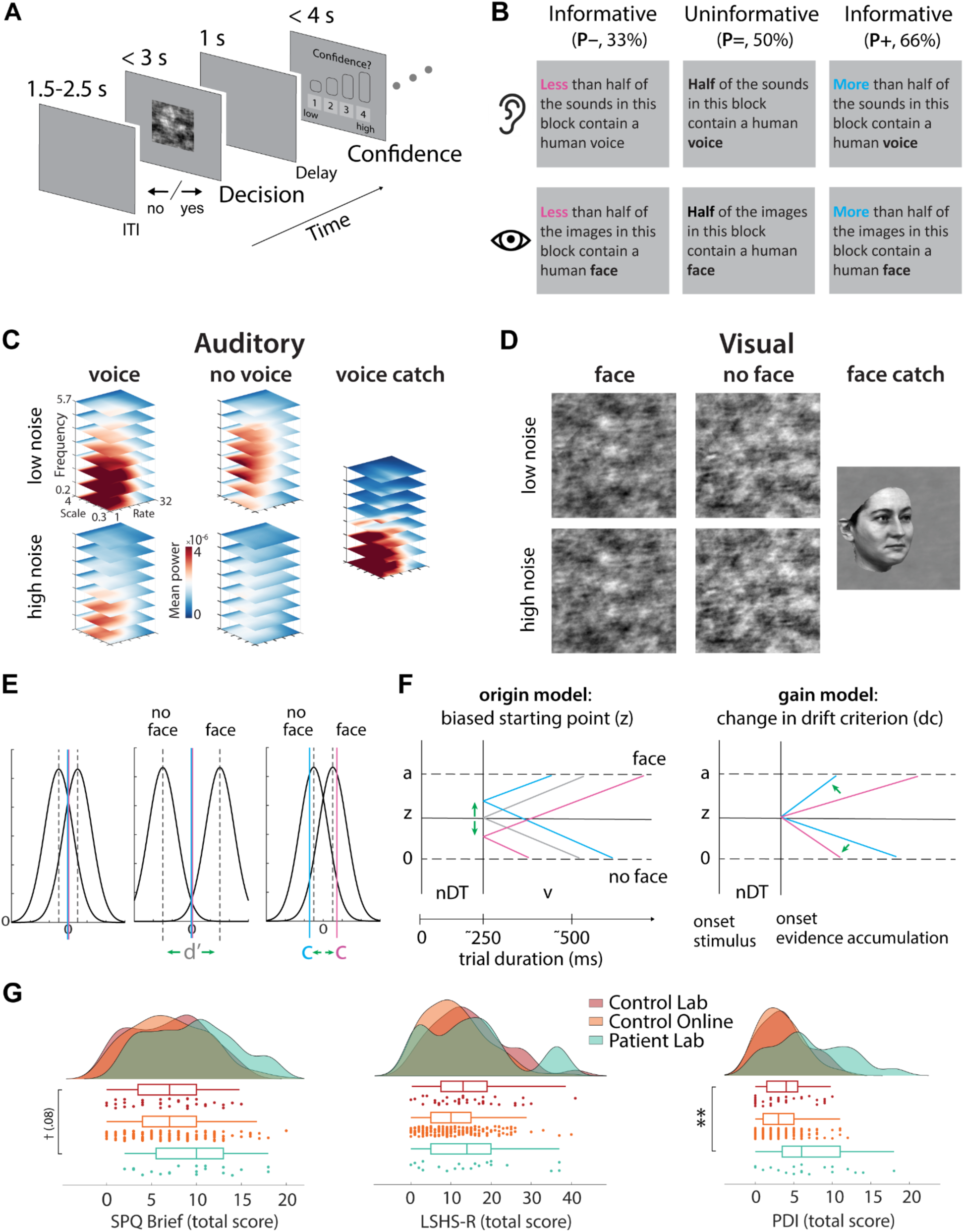
Study design, stimulus examples, state and trait information. **A)** Trial procedure (for details, see “Methods”). **B)** Manipulation of prior expectation states presented at the start of each block of 32 trials. These truthful cues indicated either an informative (less [33%] or more [66%] than half of the trials being voice or face targets) or an uninformative (50%) target probability. Target ratios were counterbalanced and randomised across the nine experimental blocks with three blocks per ratio. Only the first block was always cued by an uninformative target probability. **C)** Acoustic-energy kernels representing the average acoustic properties (cochlear frequency; temporal rate; and spectral scale) of the vocalised (left) and non-vocalised (middle) sounds as well as the average undistorted voice catch sounds (right) used in the experiment. These speech properties were generated by mixing selected statistics of the original sound and pink noise (^73^; for details, see “Methods”). The psychoacoustic power spectra of voice sounds are in line with what has previously been identified as speech-typical ^59^. The no voice high noise condition contained informative content but very little speech-typical acoustic properties. **D)** Examples of grey-scaled phase coherence modulated images of a face (left) or an image texture with the same average statistics but no specific face content (middle). A clear version of the same face image representing a potential face catch image (right). All face images were derived from the MPI face database (Max Planck Institute for Biological Cybernetics, Tübingen, Germany). The low and high noise conditions presented images at 22.5% and 17.5% phase coherence, respectively. Phase coherence of the catch images was set to 85%. **E)** Theoretical adaptations of signal detection theory (SDT; ^71^). Shift of entire signal (face) and noise (no face) distributions is depicted by thick black lines (perceptual sensitivity; d’; left). Shifts in decision bias (i.e., criterion; right) with informative prior probability cues are depicted by coloured vertical lines (pink represents a hypothetical criterion for fewer target blocks [P^−^], while the blue line represents more target blocks [P^+^]). **F)** Theoretical parameter shifts in the drift diffusion model (DDM) postulated by the origin (i.e., starting point, z) and the gain model (i.e., drift criterion, dc). Key parameters of the model are non-decision time (nDT), boundary (a), starting point (z), and drift rate (v). **G)** Raincloud plots ^74^ comparing samples with and without psychosis collected in online and laboratory environments ahead of the experiment (N_control-lab_ = 36, N_control-online_ = 192, N_patient-lab_ = 20) on three instruments examining schizotypy (SPQ Brief; ^75^, hallucination [LSHS-R; ^75,76^], and delusion [PDI; ^77^]) proneness as trait markers.

The present data show that, first, human perceivers with or without a diagnosis of psychosis all adopt a conservative decision strategy during complex auditory or visual detection tasks, particularly when sensory evidence is weak. Second, when voices or faces occur relatively rarely, prior information emphasises this overall conservative decision bias. Third, in healthy individuals, such prior information modulates a multi-stage process and especially the sensory gain during evidence accumulation. In individuals with psychosis, however, this gain is reduced, resulting in less malleable perceptual decision-making.

Our results showcase how prior information impacts strategic choice behaviour and can modulate sensory information extraction under high sensory uncertainty.

## Results

In the present study, we used derivatives of face and voice stimuli to emulate key aspects of everyday perceptual decisions. We asked which components of decision behaviour are modulated by (1) information on prior target probabilities, (2) the tested modality (auditory vs visual), (3), the amount of sensory stimulus evidence (low vs high noise). In all of this, we centrally aimed to (4) test which of these mechanisms are specifically altered in individuals with a diagnosis of psychosis.

We acquired and analysed laboratory-collected data from 20 in-patients with a psychosis diagnosis (F2x.x; ICD-10 ^78^), admitted to a psychosis ward at the university hospital Schleswig-Holstein, and data from 36 healthy age-matched participants without psychosis. Additionally, to increase statistical power of decision behaviour in healthy individuals, we present data from 192 participants without psychosis collected online via the platform Prolific (www.prolific.com). All participants took part in separate face and voice detection tasks (for details, see “Methods”; Table S1).

### Individual schizotypal traits follow a continuum

The traits schizotypy, hallucination, and delusion proneness are assumed to vary along a continuum ranging from non-clinical to clinical psychosis populations ^6,79–82^. We examined differences in these individual traits using dimensional ratings on three common self-report scales, allowing us to run analyses across healthy, subclinical and clinical expressions (Schizotypal Personality Questionnaire-Brief [SPQ-B], Launay-Slade Hallucination Scale-Revised [LSHS-R], Peter’s Delusion Inventory [PDI]; ^75–77;^ Fig. 1G).

Our obtained self-reported scores corroborate the continuum hypothesis, as participants with and without a clinical diagnosis of a psychotic disorder covered a similar range in all trait scores, with slightly higher maximum scores for participants with psychosis for delusion proneness (SPQ (max. score 22): range_control-lab_ = 0-15, range_control-online_ = 0-20, range_patient_ = 2-18; LSHS (max. score 48): range_control-lab_ = 0-41, range_control-online_ = 0-40, range_patient_ = 0-37; PDI (max. score 21): range_control-lab_ = 0-10, range_control-online_ = 0-12, range_patient_ = 0-18; Fig. 1G; Table S1). Negative binomial regression models indicated that the distributions of schizotypal traits and hallucination proneness were slightly, but not not significantly shifted between the lab control group compared to both the online control and patient group (SPQ: *β*_control-online_ = 0.022, 95% CI = [-0.22 - 0.26], *p* = .8593; *β*_patient_ = 0.315 [-0.04 - 0.67], *p* = .0824; LSHS: *β*_control-online_ = -0.269 [-0.55 - (-0.01)], *p* = .0508; *β*_patient_ = 0.035 [-0.37 - 0.45], *p* = .8669; Fig. 1G). This result provides more evidence for the continuum hypothesis. We only found delusion proneness being significantly elevated in patients with psychosis (PDI: *β*_control-online_ = -0.12 [-0.42 - 0.18], *p* = .4515; *β*_patient_ = 0.645 [0.22 - 1.08], *p* = .0032; Fig. 1G), confirming the presence of this core symptom of psychosis in our patients via self-reports ^3^.

In sum, the present data suggest a continuum of self-reported schizotypal traits that do not clearly delineate clinical from non-clinical populations.

### Informative probabilistic prior information affects decision behaviour

Flexible use of different decision strategies allows individuals to tailor their choices to environmental demands ^83^. Cues indicating target probabilities can act as exogenous decision support or bias ^20^. To assess whether participants update their expectations based on block-level probabilistic cues, we manipulated information on target probabilities at the start of each block (see “Methods”; Fig 1B).

Results from (generalised) linear mixed-effects models of our two high-uncertainty perceptual tasks, upon false discovery rate correction and accounting for antipsychotic medication, show distinct effects of prior information on decision behaviour (Tables S2-S6; for in-detail analysis of hit and false alarm rates, see Supplementary Materials Fig. S1 and Tables S3-S4).

We found a conservative decision strategy, with overall lower predicted choice probabilities than actually presented target probabilities across cues (*M* = -18.24%). In our lab data, these conservative choice probabilities generally violated parametric adaptations of choice probabilities that would have been in line with the presented target probabilities (*M*_P–_ = -8.34%, *M*_P=_ = -17.59%, *M*_P+_ = -26.49%; Fig. 2A). More specifically, while the negative informative cue (P^−^) significantly decreased target choices compared to the uninformative cue (P^=^), the positive informative cue (P^+^) did not result in a significant increase in target choices (P^−^: *OR*_lab_ = 0.64 [0.49 - 0.84], *p* = .006; P^+^: *OR*_lab_ = 1.34 [1.02 - 1.76], *p* = .075; Fig. 2A). Neither of the two informative cues interacted significantly with the modality of the task, showing supramodal modulations by prior information (P^−^: *OR*_lab_ = 1.24 [0.72 - 2.12], *p* = .593; P^+^: *OR*_lab_ = 1.40 [0.81 - 2.39], *p* = .385; Fig. 2A).

**Fig. 2.**
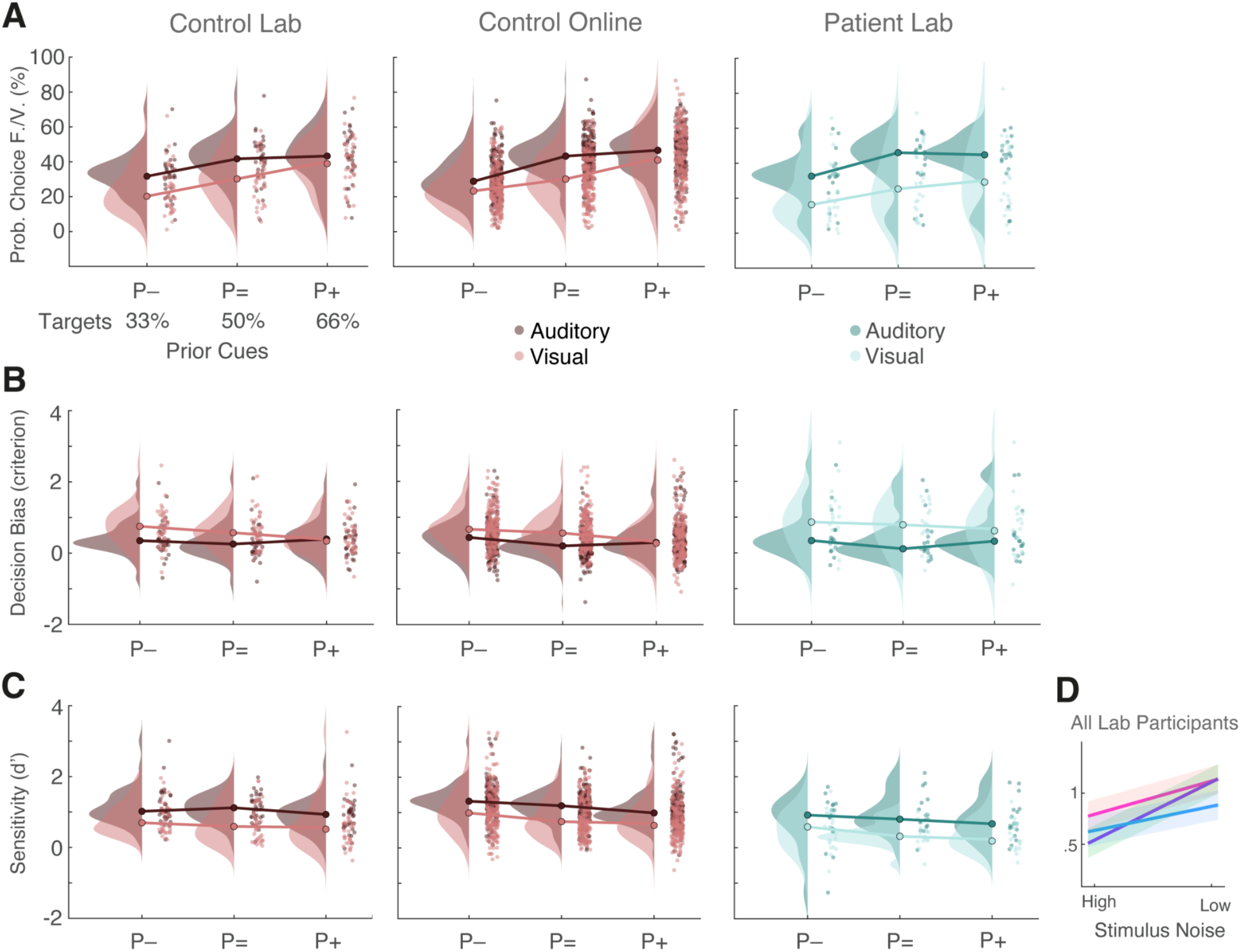
Effects of prior information on perceptual decision behaviour. Prior information cues indicated true target probabilities, presented as P^−^ (33%), P^=^ (50%), and P^+^ (66% ‘face’ or ‘voice’ probability in the current block). In panels A, B and C, reddish colours depict data of lab and online control groups (N=36, N=192, respectively), while turquoise colours depict data of patients with psychotic disorders (N=20). Dots represent average data of individual participants. Raincloud plots as a function of prior information cue, modality and group for **(A)** observed face or voice choice probabilities, **(B)** decision bias (“criterion”), **(C)** perceptual sensitivity (d’), **(D)** Predicted decision bias (“criterion”) from LMMs as a function of prior information and stimulus noise. Data collapsed across all participants collected in the lab. Blue lines depict predictions for positively (P^+^), purple lines for uninformatively (P^=^), and pink lines for negatively (P^−^) cued blocks. Data collapsed across tasks and all participants collected in the lab (N=56). High noise sounds had a mixing ratio of 50/50% (original voice/pink noise), while low noise stimuli were mixed at (95/5%). High noise images were presented at a phase coherence of 17.5% and low noise images at 22.5%.

Our online control sample without psychosis corroborated the prior-specific significant decrease in choice probabilities with the negative informative cue (P^−^: *OR*_online_ = 0.54 [0.39 - 0.75], *p* = .001; Fig. 2A). Additionally, these well-powered online control group also showed a significant increase in target choices with the P^+^ cue (*OR*_online_ = 1.45 [1.05 - 2.01], *p* = .049; Fig. 2A). Despite this significant increase, choice probabilities with the P^+^ cue remained well below the respective target probabilities, underlining the overall conservative decision behaviour (*M*_all_cues_ = -18.24% vs *M*_P+_ = - 26.49%). These results also indicate a smaller modulatory effect of the P^+^ cue among informative cues, whose detection necessitates a large dataset, as underlined by our online sample. Notably, people with and without psychosis did not differ significantly in their overall choice probabilities (*OR*_main_effect_ = 0.46 [0.23 - 0.93], *p* = .075; Fig. 2A). Neither did any prior information cue affect choice probabilities differentially across tasks in patients with psychosis (P^−^: *OR*_patient_interaction_ = 1.13 [0.96 - 1.33], *p* = .314; P^+^: *OR*_patient_interaction_ = 0.93 [0.79 - 1.09], *p* = .502; Fig. 2A).

Overall, while all participants adapted their choices in accordance with the presented cues, the magnitude of this adaptation varies between informative cues. The inconsistent adaptation of choice probabilities in line with target probabilities across datasets, specifically in P^+^ cued blocks, indicates a more conservative strategy as a consequence of more cautious strategic adaptations than with the two other cues.

To provide more comprehensive insights into choice behaviour as a function of decision performance and bias, we examined potential changes in perceptual discriminability (i.e., sensitivity) and decision bias (i.e., criterion) with prior information. To this end, we used linear mixed-effects models predicting the two main parameters from SDT sensitivity and criterion ^71^.

The manipulation of prior information on target probabilities should, according to SDT postulations, elicit strategic shifts in decision bias (“criterion”; ^83,84^). These shifts could be set a priori at the start of a block of trials, as used here. In line with these postulations, we found that our manipulation of prior information results in several changes in decision bias. Specifically, compared to the uninformative cue (P^=^), we observed a general significant increase in decision bias in P^−^ cued blocks across all participants in the lab data (P^−^: *β*_lab_ = 0.09 [0.02 - 0.16], *p* = .024; Fig. 2B). This is complemented by a significant general reduction in bias in P^+^ cued blocks in our online data (P^+^: *β*_online_ = -0.26 [-0.30 - (-0.22)], *p* < .001; Fig. 2B). These differential effects can be separated further by taking task modality into consideration, as decision bias was generally higher in the visual task across both datasets (*β*_lab_ = 0.32 [0.17 - 0.46], *p* < .001; *β*_online_ = 0.26 [0.17 - 0.35], *p* < .001; Fig. 2B). Though, this bias increase is of smaller magnitude in P^+^ cued blocks in the lab data, resulting in the P^+^ cue significantly interacting with modality in these data (P^+^: *β*_lab_ = -0.26 [-0.39 - (-0.12)], *p* = .001; Fig. 2B). In other words, a small increase in decision bias in visual P^+^ cued blocks is contrasted by a stronger increase in both visual P^−^ and P^=^ cued blocks. Again, we did not find evidence for patients with psychotic disorders adapting their decision strategy (i.e., bias) differently across prior information cues (*β*_lab_ = 0.44 [0.01 - 0.87], *p* = .102; Fig. 2B). Hence, the observed differences in decision bias also suggest prior information-specific adaptations of decision criteria across participants that could be strategic and conservative in nature with the aim of avoiding false target detections in blocks with more non-target stimuli.

Complementarily, we examined whether the decision strategy chosen in accordance with prior information cues benefitted performance, as indicated by perceptual sensitivity. The results demonstrate beneficial effects of the negatively informative cue (P^−^) on decision performance. That is, perceptual sensitivity increased in P^−^ cued blocks across all participants in the lab data (P^−^: *β*_lab_ = 0.13 [0.02 - 0.24], *p* = .050; Fig. 2C). Our online control sample allowed us to differentiate the effects of the P^−^ cue further. Here, the P^−^ cue also interacted significantly with modality, resulting in the highest sensitivity among cues in the auditory task but a low sensitivity in the visual task (P^−^ : *β*_online_ = -0.18 [-0.31 - (-0.04)], *p* = .015; Fig. 2C).

Contrarily, the informative positive cue (P^+^) did not benefit performance, as evidenced by significantly decreased sensitivity in our online control group (P^+^: *β*_online_ = -0.09 [-0.15 - (-0.02)], *p* = .015; Fig. 2C).

Similar to our results on choice probabilities, people with and without psychotic disorders did not differ significantly in their overall sensitivity after controlling for antipsychotic medication (*β* = -0.22 [-0.51 - 0.06], *p* = .186) nor were patients’ sensitivity affected differently by any prior information cue (P^−^: *β* = -0.16 [-0.40 - (-0.07)], *p* = .254; P^+^: *β* = 0.04 [-0.20 - 0.28], *p* = .800; Fig. 2C).

We further examined the effects of stimulus noise, which is a crucial determinant for decision behaviour under uncertainty ^11^, and informative cues on SDT parameters. Our results showed that both informative cues interacted significantly with stimulus noise across tasks and groups in the lab (P^−^: *β*_lab_ = -0.27 [-0.49 - (-0.05)], *p* = .049; P^+^: *β*_lab_ = -0.37 [-0.59 - (-0.14)], *p* = .005; Fig. 2D). More specifically, for more uncertain stimuli (i.e., high noise) both informative cues increased sensitivity compared to the uninformative cue. However, for less uncertain stimuli (i.e., low noise), no informative cue (P^−^/P^+^) benefitted sensitivity—with the P^+^ cue even resulting in slightly reduced sensitivity (Fig. 2D).

To summarise, these results demonstrate that all participants adapted their choice behaviour in our perceptual decision tasks in response to block-by-block varying probabilistic prior information on target probabilities strategically and independently of psychotic disorders. Prior information can thus benefit perceptual decision outcomes—particularly under high uncertainty. However, participants did not use both informative cues symmetrically. That is, although being equally informative as its P^−^ counterpart, the P^+^ cue appeared to be generally less beneficial for decision behaviour and performance; especially so in the visual condition. Our finding that the informative cues’ utility increases with increasing stimulus noise suggests that prior information builds an internal reference state for a block of trials, which is subsequently modulated by the incoming amount of stimulus noise (i.e., sensory evidence) on a trial-by-trial basis.

### Schizotypal traits and psychotic symptom strength modulate sensitivity and bias

Prior information affects only a few decision behaviour parameters and self-reported traits significantly in patients with psychotic disorders. These findings suggest no clear dichotomous boundaries between clinical and non-clinical populations ^85^ (Fig. 1G). Therefore, we sought to examine associations between the examined traits (i.e., schizotypy, hallucination and delusion proneness) and decision performance metrics across all participants of an environment.

The lab-collected data did not show significant effects of any self-reported trait on perceptual sensitivity or decision bias (Tables S7a, S8a). Schizotypy did not significantly interact with clinical psychosis diagnosis as predictors of sensitivity, providing more evidence for a continuum (*β*_interaction_lab_ = 0.06, CI = [-0.02 - 0.13], *p* = .329). However, in the online data, we observed a schizotypy-by-modality interaction: sensitivity decreased with higher proneness to schizotypy in the visual task, whereas schizotypy did not have a general effect on sensitivity in the auditory task (*β*_interaction_online_ = -0.03 [-0.05 - (-0.01)], *p* = .036; Table S7b; Fig. 3A). This interaction could be differentiated further by adding prior information as a third factor; resulting in linking higher schizotypy proneness to increased perceptual sensitivity in P^−^ cued blocks in the auditory task (*β*_interaction_online_ = -0.04 [-0.06 - (-0.02)], *p* < .001; Fig. 3A). Contrarily, higher schizotypy proneness reduced sensitivity with both informative cues (P^−^/P^+^) in the visual task (*β*_interaction_online_ = -0.02 [-0.04 - (-0.01)], *p* = .041; Fig. 3A). The uninformative P^=^ cue showed no variation with schizotypy proneness in either task. Hence, schizotypy—a general trait—and prior information affected participants’ perceptual signal-to-noise ratio in the two modalities in opposite directions.

**Fig. 3.**
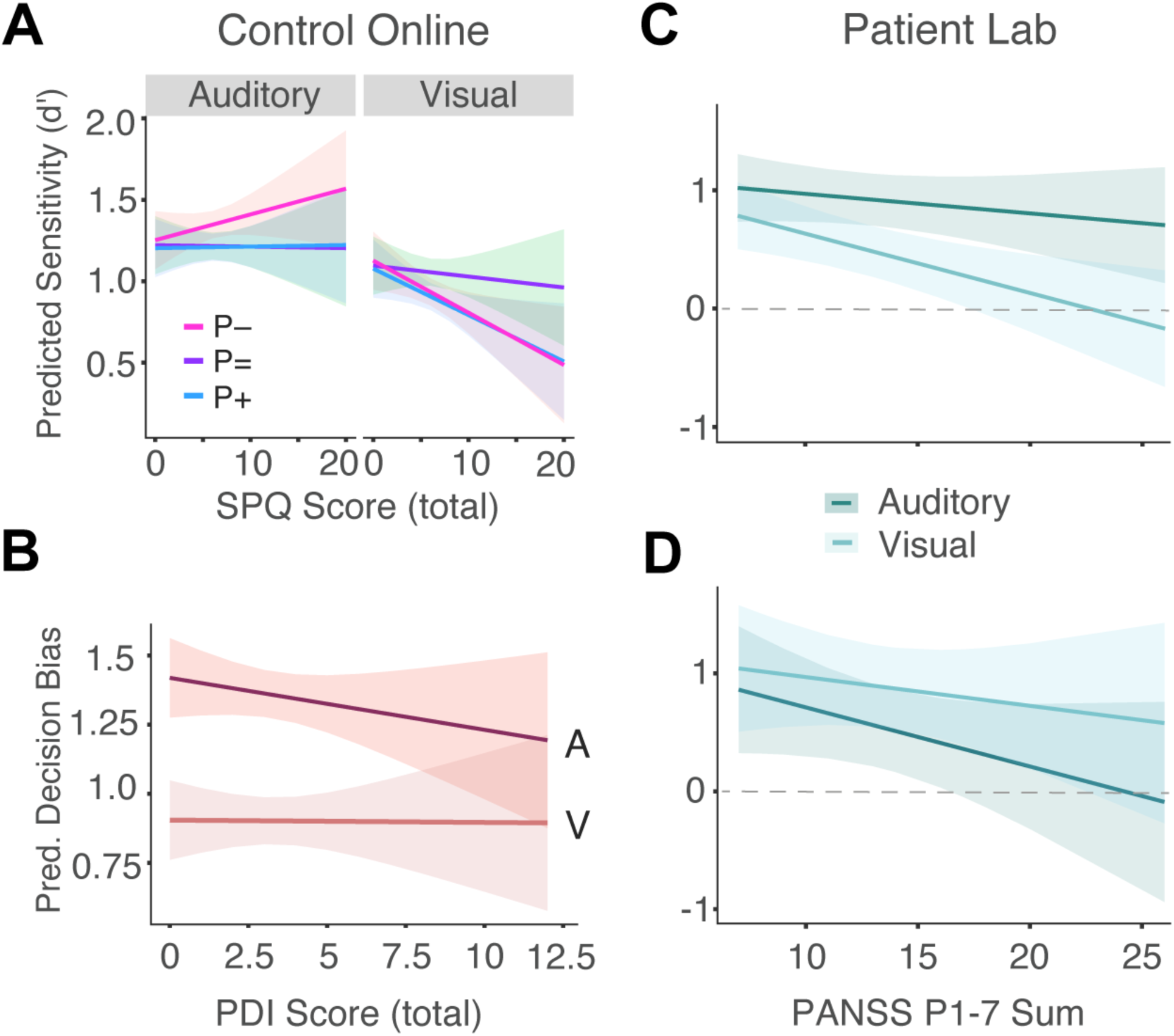
Personality traits and positive symptoms are linked to decision performance. **A)** Predicted perceptual sensitivity (d’) from a LMM as a function of total SPQ score ^75^ and modality for the online control group without psychotic disorders. Blue lines depict predictions for positively (P^+^) cued blocks, purple lines for uninformatively (P^=^) cued blocks and pink lines for negatively (P^−^) cued blocks. **B)** Predicted decision bias (“criterion”) as a function of total PDI score ^77^ and modality for the online control group. **C)** Predicted perceptual sensitivity (d’) from a LMM as a function of the sum of positive symptoms around the time of testing and modality for patients with psychosis who took part in the lab. Darker colours depict estimates for the auditory task, while lighter colours depict estimates for the visual task. **D)** Predicted decision bias as a function of the sum of positive symptoms around the time of testing and modality. Same group and colour scheme as in panel C.

For decision bias, a different trait interacted with task modality in the online data. Here, delusion proneness negatively correlated with decision bias in the auditory task, while this relationship was absent in the visual task (*β*_interaction_online_ = -0.03 [0.01 - 0.05)], *p* = .007; Table S8b; Fig. 3B). We found no significant predictive power of any other self-reported trait, and combinations with prior information, on decision bias.

The previous analyses of decision behaviour varying with general personality traits underline the importance of considering relevant traits of the decision maker. These analyses have also demonstrated that proneness to traits linked to psychosis (i.e., schizotypy) and positive symptoms (i.e., delusions) systematically affect decision-making performance across non-clinical populations. In addition, since self-report questionnaires may not capture the symptomatology of patients with psychosis around the time of testing accurately, we obtained more detailed ratings of positive, negative and general symptoms from all patients with psychosis right after testing by conducting systematic PANSS interviews ^86^.

We found a negative association of both sensitivity and decision bias with the sum of positive symptoms, which is independent of prior information. Though, this association interacted with modality again (*β*_interaction_lab_ = -0.03 [-0.05 - (-0.02)], *p* < .001; Tables S9-S10; Fig. 3C, D). For sensitivity, this negative association was stronger in the visual modality, while, for decision bias, it was stronger in the auditory modality (Fig. 3C, D). These associations resemble the ones observed across all participants. Their direction remained the same, although not significant, when examining effects by prior information separately.

Taken together, prior information and proneness to schizotypal personality, a non-clinical construct of schizophrenia, affect how stimuli are perceived in auditory and visual perceptual decision-making. Additionally, higher proneness to delusions may result in adopting more liberal decision criteria, at least in the auditory task. Similarly, patients with psychotic disorders who experience more severe positive symptoms around the time of testing apply a more liberal decision strategy, resulting in worse perception across both modalities. These links between perceptual decision performance and the severity of positive symptoms underline the perceptual challenges of psychotic disorders and potential compensatory bias mechanisms captured by our real-world inspired target detection tasks.

### Participants trust exogenous prior information

Upon completing the experiment, all participants rated their trust in the validity of our prior information cues on a 5-point Likert scale (dishonest to honest). The trust ratings of all groups are generally in the mid-range across tasks (*M*_control-lab_ = 2.43, *SD*_control-lab_ = 1.03; *M*_control-online_ = 3.08, *SD* = 1.24; *M*_patient_ = 3.09, *SD* = 1.07). An ordinal mixed regression model showed significantly less self-reported trust only for lab control participants compared to online control participants (*β* = - 1.78, *SE* = 0.051, *p* < .001). All participants believed in the truthfulness of the provided information with no significant difference between modalities (*β* = -0.14, *SE* = 0.269, *p* = .6013; BF = 0.46).

### Prior information and psychosis modulate sensory gain

Signal-detection measures lack temporally resolved information on how much individual components, including different forms of bias, contribute to the final choice ^87^. Therefore, we next investigated decision performance using single-trial response times (RTs). RTs are indicative of the amount of cognitive processing required to do a task and represent a continuous proxy of decision support prior information may provide. Therefore, it is conceivable that prior information affects target choices in faster and slower trials differently ^20^.

In single-trial response times, linear mixed modelling revealed general and modality-specific effects of prior information, while controlling for antipsychotic medication (Tables S11). Specifically, decisions were generally fastest in P^−^ cued blocks in both our lab and online data (*β*_lab_ = -0.03, CI = [-0.04 - (-0.01)], *p* = .001; *β*_online_ = -0.04 [-0.05 - (-0.03)], *p* < .001; Fig.. 4A). In our healthy online sample, mixed modelling showed generally faster RTs in P^+^ cued blocks as well (*β*_online_ = -0.04 [-0.05 - (-0.02], *p* < .001; Fig. 4A). Further, in the online data, we observed that the P^+^ cue supports the decision process by speeding up decisions specifically in the auditory but not the visual task (*β*_interaction_online_ = 0.05 [0.02 - 0.08], *p* = .001; Fig. 4A). Patients with psychosis benefited less from the P^−^ cue than their counterparts without psychosis (*β*_interaction_lab_ = 0.03 [0.01 - 0.05], *p* = .013; Fig. 4A, B), particularly on fast trials (Fig. 4B). Hence, these results demonstrate faster decisions with informative cues across datasets but a smaller benefit for patients with psychosis that is specific to the negative informative cue. They further suggest an initial bias of the decision process, which could be captured by the most prominent model of decision-making, the DDM ^26^.

**Fig. 4.**
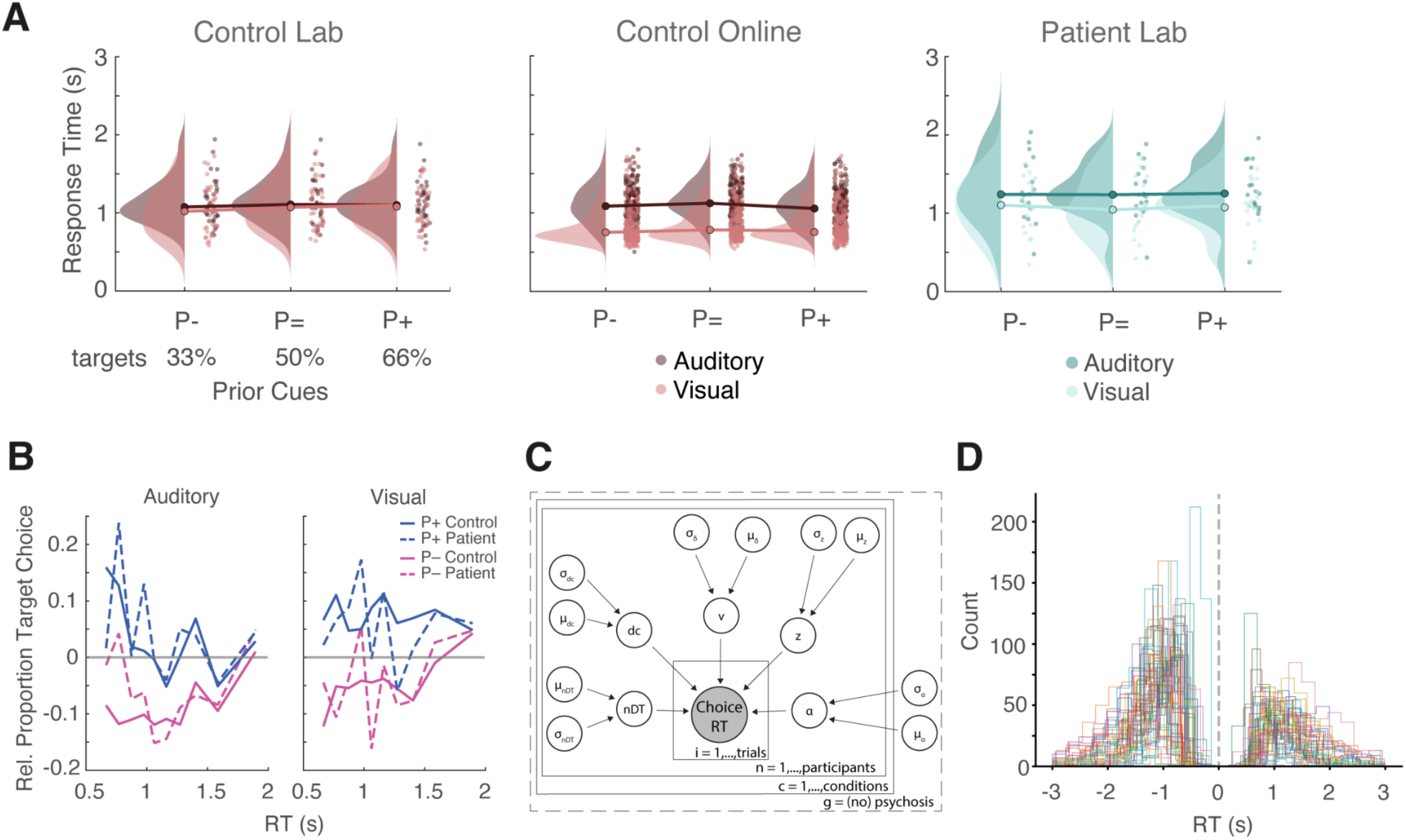
Modelling shows aberrant response times in psychotic disorders. **A)** Raincloud plots of observed median response time in seconds as a function of prior information, modality, and psychosis diagnosis. **B)** Cumulative response functions (CRF) by informative prior cues and psychosis diagnosis. Lines represent the proportion of target choices of each informative cue (P^−^/P^+^) relative to the uninformative P^=^ cue (baseline) for each decile of the respective RT distributions. Solid grey lines represent no change in target choice proportion with an informative cue. Dashed blue and pink lines depict data from participants with a clinical psychosis diagnosis, while their solid counterparts depict data from lab control participants. **C)** Model diagram of two identical stimulus-coded HDDMs outlining fitted parameters (nDt, z, v, a, dc) based on RT distributions as a function of choice (i.e., target vs non-target choice). **D)** Individual participant RT distributions (in seconds) split by choice (negative RTs indicate non-target choices) across all lab participants. Vertical dashed grey line represents stimulus onset. This flipped representation was used in the drift diffusion modelling.

As a final step on our analysis, we employed hierarchical drift-diffusion models (HDDM) to warrant a more mechanistic examination of differential effects of probabilistic prior information on the decision-making process.

In this model, prior information could 1) shift the evidence accumulation starting point (origin model; ^13,88^) or 2) bias the evidence accumulation itself (gain model; ^13,89^). The starting point (*z*) reflects an initial predisposition toward one decision, while the drift criterion (*dc*) reflects biases in the interpretation of ambiguous stimulus evidence during evidence accumulation ^32^. To disentangle modulations of these two decision parameters, we employed a hypothesis-driven, stimulus-coded hierarchical drift diffusion model (HDDM; ^26,57,58,90,91^; Fig. 4C, 4D; see “Methods”). To benefit from hierarchical dependencies without confounding results by mixing groups, we ran identical models separately for lab participants with and without psychotic disorders (Fig. 4C).

Our HDDM results reveal that multiple components of the decision process are affected during audio-visual decision-making under high uncertainty. General differences in modality-specific temporal stimulus dynamics, as expected, act on the sensory encoding phase, since we found longer non-decision times (nDT) in the auditory task across participants (control: *M*_auditory_ = 0.658 s, 95% HDI [0.604, 0.711], *M*_visual_ = 0.597, 95% HDI [0.543, 0.652]; patient: *M*_auditory_ = 0.610, 95% HDI [0.498, 0.712], *M*_visual_ = 0.527, 95% CI [0.427, 0.622]; P_control_[auditory posterior distribution > *M* visual posterior distribution] = 0.94; P_patient_[auditory > *M* visual] = 0.89; Fig. 5A). Participants with psychosis showed slightly shorter nDT across modalities (P[control > *M* patient auditory] = 0.79; P[control > *M* patient visual] = 0.89; Fig. 5A). These results suggest that all participants take longer for sampling auditory evidence, which unfolds over time instead of being static, with slight differences in psychosis.

**Fig. 5.**
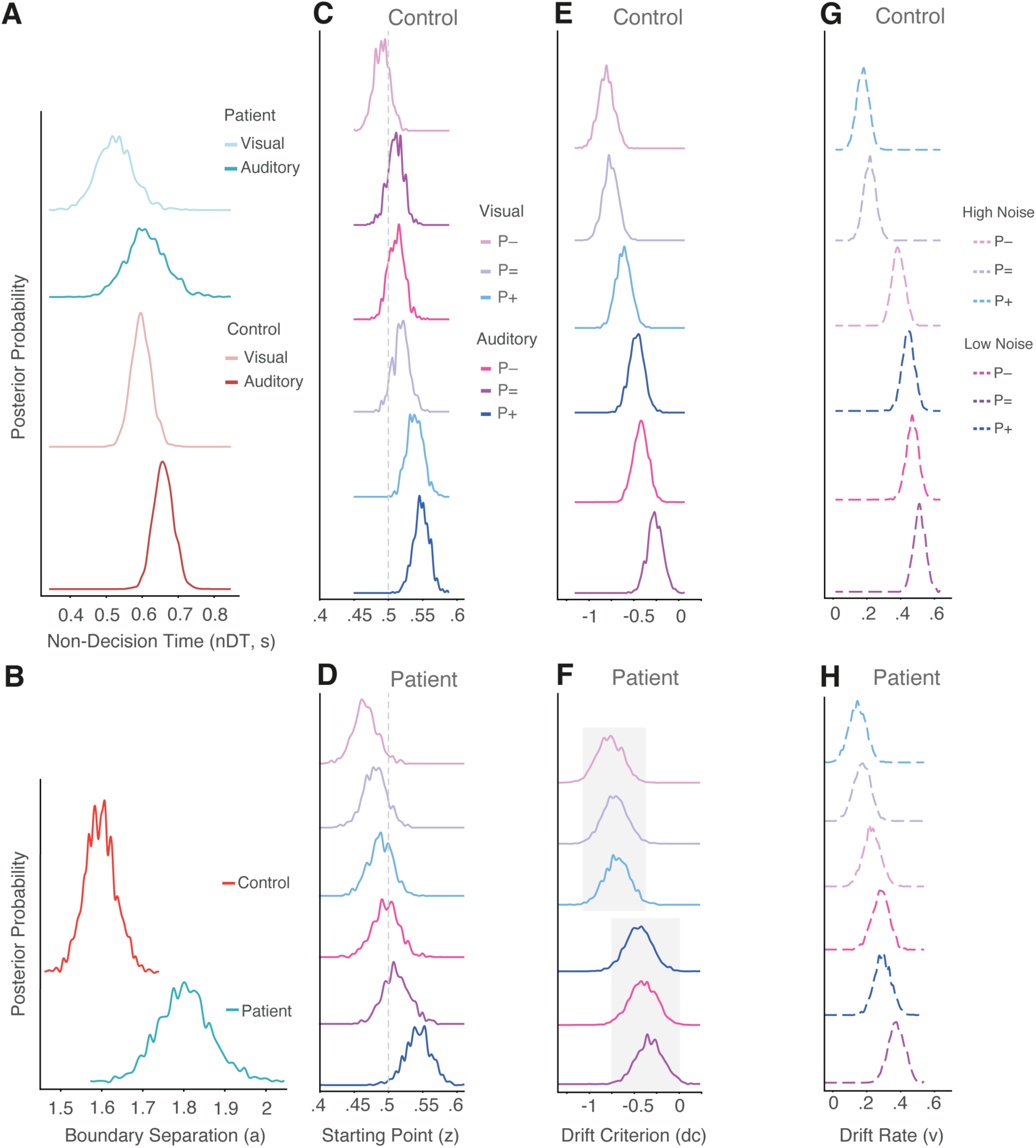
Modelling of response times and choice shows aberrant evidence accumulation in psychosis. Raincloud plots of observed median response time in seconds as a function of prior information, modality, and psychosis diagnosis. **A)** Comparison of posterior probability estimates of non-decision time (nDT, in seconds) as a function of modality and psychosis. Reddish colours depict data from participants without psychosis, while turquoise colours depict estimates for participants with psychosis. Darker shades depict estimates for the auditory task, while lighter colours depict visual task estimates. **B)** Comparison of posterior probability estimates of boundary separation (a, a.u.) as a function of psychosis. **C, D)** Posterior probability estimates of the starting point (z) of evidence accumulation as a function of prior information and modality. Blue distributions depict predictions for positively (P^+^), purple lines for uninformatively (P^=^), and pink lines for negatively (P^−^) cued blocks. Lighter shades depict visual task estimates, while darker shades depict estimates for the auditory task. While panel C depicts data from the lab control group, D depicts data from patients with psychotic disorders. **E, F)** Posterior probability estimates of the drift criterion (dc) as a function of prior information and modality. Colour code and organisation as in panels C and D. **G, H)** Posterior probability estimates of the drift rate (v) as a function of prior information and stimulus noise. Lighter shades depict estimates for trials with high stimulus noise, while darker shades depict estimates for trials with low stimulus noise. Organisation as in panels F and G.

Further, completely separate highest density intervals (HDIs) provided evidence for an increased decision boundary separation in participants with psychosis (*M*_control_ = 1.60, 95% HDI [1.525, 1.674], *M*_patient_ = 1.80, 95% HDI [1.686, 1.948]; P(control > *M* patient) = 0.997; Fig. 5B). This difference of posterior distributions indicates that, in general, patients with psychosis require more evidence to commit to a choice and prioritise accuracy over response speed.

Prior information modulated several components of the decision process representing biases, but did so to different extents in individuals with and without psychosis. First, we observed changes in the starting point (*z*) with informative cues mostly in line with target probabilities, which cover a wider range in patients with psychosis (*M*_control_ range = 0.491-0.547, *M*_patient_ range = 0.467-0.542; Table S12; Fig. 5C, D). Across participants, these shifts indicating an a priori bias or expectation, were most prominent for the informative P^+^ cue (control: P(P^+^_auditory_ > 0.5) = 1, 95% HDI [0.525, 0.569], P(P^+^_visual_ > 0.5) = 1, 95% HDI [0.516, 0.562]; patients: P(P^+^_auditory_ > 0.5) = 0.983, 95% HDI [0.506, 0.579], P(P^+^_visual_ > 0.5) = 0.269, 95% HDI [0.455, 0.524]), but were also present for the visual P^−^ cue (control: P(P^−^_visual_ < 0.5) = 0.78, 95% HDI [0.471, 0.516]; patients: P(P^−^_visual_ < 0.5) = 0.965, 95% HDI [0.434, 0.506]; Fig. 5C, D). Particularly in the group without psychosis, the P^+^ cue led to a supramodal starting point shift—in line with our SDT results on decision bias (Fig. 2B). Interestingly, patients with psychotic disorders shifted their starting point less towards target choices with the P^+^ cue in the visual task (*M*_visual_: P^+^_control_ = 0.538, 95% HDI [0.516, 0.562], P^+^_patient_ = 0.489, 95% HDI [0.455, 0.524]; Fig. 5D), suggestive of conservative choice behaviour.

Most interestingly, we found the overall conservative decision strategy resulting in a negative drift bias (dc) across all conditions and participants (Table S13; Fig. 5E, F). This evidence-processing bias indicates that all participants interpret the provided ambiguous evidence as supporting non-target choices. However, only individuals without psychosis amplified this interpretation of the evidence accumulation process with prior information, in line with the gain model ^13^. Differences in this gain followed target probabilities (i.e., P^−^ < P^=^ > P^+^) in the visual task, while both informative cues increased drift bias more negatively in the auditory task (P^−^/P^+^ < P^=^; Table S13; Fig. 5E, F). The variation in drift bias with prior information was considerably reduced in patients with psychosis (i.e., more overlap between cue distributions, P(minimum overlap): V_control_ = 0.063, A_control_ = 0.046; V_patient_ = 0.34, A_patient_ = 0.276; Fig. 5F). These findings strongly indicate that the evidence accumulation process is more prominently modulated by probabilistic cues due to implicit expectations that take effect during a trial in healthy individuals.

Stimulus noise, in line with perceptual decision-making theory ^11^, modulated the quality of the evidence accumulation (drift rate, *v*). That is, participants accumulated evidence faster in trials with low stimulus noise (low noise: *M*_control_ range = 0.501–0.567, *M*_patient_ range = 0.294–0.373; high noise: *M*_control_ range = 0.237–0.443, *M*_patient_ range = 0.148–0.231; Fig. 5G, H). Though, this low-noise benefit was generally reduced in patients with psychosis who showed slower drift rates across both prior information and stimulus noise conditions (P(P^=^_control_ > *M* P^=^_patient_) = 0.999, P(P^+^_control_ > *M* P^+^_patient_) = 0.929; Fig. 5G, H).

Taken together, our computational modelling results demonstrate that probabilistic prior information induces separable biases in perceptual decision-making under high uncertainty, with differences between individuals with and without psychotic disorders in when these biases influence the decision process. Specifically, while we find a modulation of the sensory gain allowing for the incoming stimulus evidence to be interpreted more efficiently by individuals without psychosis, patients with a clinical psychosis diagnosis rather change their a priori expectation towards the stimuli and shorten their sensory encoding time. Individuals without psychosis also make use of shifting their a priori expectation with target probabilities, but to a smaller extent.

## Discussion

Exploiting probabilistic information allows us to adapt our decisions to environmental demands and can act as an exogenous decision support, or bias ^20,92^. The present study investigated how exogenous probabilistic prior information affects perceptual decision-making under high sensory uncertainty across the auditory and visual modalities in individuals with and without psychotic disorders.

Our study aimed to shed light on whether prior information supports the dynamic and multifaceted decision process ^11^ a priori (origin model; ^13^), during the interpretation of incoming sensory evidence (gain model; ^13^) or both proportionally (multi-stage model; ^35,36^). Since previous studies differ considerably in their task design, complexity, requirements and modalities, evidence put forward for both mechanisms is unsurprising (e.g., ^19,20,24,35,36,93^).

Here, in behavioural data, we demonstrate that probabilistic cues affect a distinct, multistage interplay of a priori bias and biased evidence interpretation across auditory and visual modalities in healthy individuals without a psychotic disorder. This group predominantly amplifies incoming sensory evidence and its accumulation speed in line with prior information. Contrarily, patients with psychotic disorders predominantly modulate a priori biases that act before encountering sensory evidence, indicating a reduced ability to exploit prior information dynamically to facilitate moment-to-moment information extraction (Fig. 5C-F).

These computational findings of a dynamic use of probabilistic information in healthy individuals provides new mechanistic support for dynamic modulations of perceptual decision components in noisy real-world environments, akin to the multi-stage model ^35^. Participants carried individual a priori expectations forward into a block of trials, where these modulate the short-term decision prior that acts as a reference for the incoming sensory evidence ^35,36^. This prepares the individual to react flexibly to variations in new, subsequent stimuli as they occur inherently in natural environments.

The advantage of a mechanistic computational modelling approach, providing temporally resolved insights into the decision-making constituents, is further underlined by a partial mismatch between our findings and SDT predictions. Recent research underlines this advantage in explaining cognitive decision-making processes ^27^. Evidence from simple perceptual tasks suggests that probabilistic cues affect the decision criterion (i.e., bias) but not perceptual sensitivity by facilitating decision performance through a top-down strategy adaptation ^13,20,23,83,84^. Here, we find probabilistic cues modulating both decision criterion and sensitivity, but to different extents. This discrepancy may simply arise from considerable variation in the employed paradigms and task/stimulus complexity between studies ^94^, where increasing complexity might result in less clear separations between bias and perceptual sensitivity parameters than SDT theory would predict. Paradigms using trial-based cues in simple perceptual tasks (e.g., ^23,41,45,51^) may not be able to play on the outlined multi-stage mechanisms because the expectation for incoming sensory evidence is being reset at the beginning of each trial just milliseconds before encountering the sensory evidence and not strategically a priori. These differences in paradigms could also result in perception aligning more with a bias on perceptual history or external sensory information during sensory analysis, which have been linked to increased vs decreased decision biases, respectively ^95^. Hence, with increasing uncertainty, efficient processing of sensory evidence appears to require a more complex adaptation of decision behaviour than simply adapting only one of the two decision processes—either one’s strategy prior to encountering a stimulus (origin model) or amplifying the processing of incoming evidence (gain model).

These crucial experimental differences show that prior information modulations in healthy decision-makers predominantly increase the gain of incoming sensory evidence. This provides more evidence for facilitatory effects in the evidence accumulation process of decision formation that behavioural and neural evidence has recently been mounting for ^28,30,34–39,93,96,97^. A conceivable neural mechanism is the strengthening of feedforward and feedback connectivity between sensory and prefrontal cortex ^93^ rather than a bias of the baseline activity of feature selective sensory units ^21,23^.

However, identifying the distinct interplay of a priori and evidence accumulation biases has been difficult and studies reporting drift biases are particularly scarce ^24^. Though, the latter has emerged as a crucial parameter when investigating perceptual decision formation ^29,32,35,36^. The fact that we observed both starting point and drift criterion modulations in healthy observers underlines the difficulty in separating these two biases clearly. Perceptual decision formation under high uncertainty may inherently prevent a full dissociation of these biases based on behavioural data alone ^24,30^. Nevertheless, our data demonstrates clear differences between participants with and without psychosis. In line with the strong prior hypothesis ^6,64^, the patient group showed virtually no variation in the drift criterion due to probabilistic information, while the healthy group showed clear modulations of this bias parameter. Support for these differences in psychosis comes from an uncertainty task showing a failure to update inference information after the introduction of new evidence at change points ^9^. These results showcase our computational model’s ability to 1) simultaneously quantify the influence of both potentially biased decision processes (origin and gain) and 2) dissociate healthy from aberrant perceptual decision-making.

We also show that healthy decision makers amplify their evidence accumulation speed, especially under higher and less predictable sensory uncertainty (Fig. 5G). This modulation of drift rate and perceptual sensitivity with sensory noise corroborates the idea that high uncertainty can be flexibly accommodated by a trial-based adaptation process that is captured by sensory analysis and evidence accumulation speed ^30,31,98^. Particularly, the P^−^ cue benefits sensitivity and evidence accumulation speed in high noise trials (i.e., weaker signals). This amplification of sensitivity and decision integration bias is further supported by previous experimental work ^23^ and by theoretical accounts of optimal decision-making in difficult-heterogeneous environments ^99^. Simultaneously, it speaks against reports of variation in stimulus uncertainty across trials not driving this effect ^38^. Hence, this finding indicates that informative probabilistic cues reduce the perceptual challenges posed by noise across modalities, reinforcing a Bayesian view of perception ^2,100^.

Notably, across all levels of analyses, we find asymmetric informative cue effects. Participants showed a particularly conservative decision (i.e., target detection) strategy with the P^+^ cue, even though it was equally informative as its P^−^ counterpart. This conservative strategy has previously been linked to changes in starting points ^18,101^. Here, we first find a lack of an ideal increase of choice probabilities in line with target probabilities for the P^+^ cue across all participants. Secondly, since fastest responses affect the DDM starting point the most, significant supramodal shifts of the respective starting points indicate that healthy individuals seem to form explicit face/voice (P^+^) predictions a priori that result in particularly fast target choices with this cue (Fig. 4B, 5C). However, this cues predictions (and benefit) seem to be less strong for slower responses, resulting in fewer target choices than expected on those slow-response trials, which could in turn drive this cue’s conservative decision behaviour.

Generally, a conservative underweighting of prior information, compared to optimal predictions from Bayesian integration models ^2,12^, has been attributed to unclear reliability of or imprecision in the prior in perceptual decision formation ^102,103^. It is conceivable that our tasks’ high uncertainty levels that fluctuated from trial to trial have made target detection difficult even after sampling more sensory evidence and resulted in participants opting for an overall greater bias for the subjectively more “reliable” stimulus category (here, no face/voice category). This is illustrated by higher decision bias (i.e., criterion) in P^−^ cued blocks, particularly in the visual task, which participants also performed worse in. Thus, a conservative decision strategy might dominate in healthy decision-making under high uncertainty to avoid false alarms and reduce the costs of decision errors.

Detection studies on prior information in clinical psychosis are scarce. Our novel approach combined auditory and visual detection tasks with probabilistic manipulations. The present examination of mechanisms of prior information proved valuable in pinpointing a failure to capitalise on the dynamic interplay of decision biases as a mechanism likely driving aberrant perception in psychosis. instead relying solely on starting point shifts. The importance of exploiting probabilistic information for neural gain during evidence accumulation itself is underlined by healthy decision makers ^30^. Psychosis is here seen to leave this stage unmodulated, as also indicated by reduced gain with lower stimulus noise levels. This aligns with previous findings on non-perceptual decision-making in clinical psychosis showing reduced cognitive processing speed and control ^52,53^.

This key difference in perceptual decision formation was primarily captured by RTs as a function of choice behaviour. SDT measures showed no general effects of psychosis or differential use of prior information after controlling for medication. The lack of differences in perceptual sensitivity contrasts with prior reports of face detection and speech-in-noise difficulties in schizophrenia ^104–108^ but finds backing in SDT predictions postulating probabilistic prior information eliciting criterion shifts. While a stronger decision bias could compensate for sensory processing imprecision, SDT criterion showed no general differences in psychosis either. Yet again, these results point to the limitations of SDT-based measures when capturing trial-to-trial cognitive variations in decision-making.

Our dimensional analysis linked perceptual sensitivity and decision bias reductions to patients’ positive symptomatology (PANSS). More severe symptoms correlated with a liberal bias in the auditory task, aligning with prior findings in voice hearers ^107^. A liberal bias in auditory tasks may reflect the auditory system’s evolutionary predisposition toward speech perception ^8,49^. If patients with more severe positive symptoms attribute aberrant salience to incoming sensory information potentially conveying speech, which is not effectively constrained by reliable cognitive or compensatory priors ^4,65^, this misattribution may exacerbate an evolutionary predisposition in the auditory domain. This interpretation gains in relevance due to higher delusional ideation being linked to a liberal bias across all participants only in our auditory task.

While SDT metrics hinted at altered expectations, RTs showed that patients benefited less from informative cues, especially in the fastest auditory and medium-fast visual trials, indicating reduced adaptive flexibility under varying uncertainty ^109^. Notably, the examined patients with psychosis failed to utilise exogenous prior information to interpret ambiguous sensory information. This overall reduced modulation of evidence accumulation suggests an overreliance on endogenous priors (origin model), thought to underlie established psychosis ^4^. Further, the identified impaired combination of endogenous priors and sensory information mirrors findings in hallucination-prone healthy individuals ^110^.

Methodologically, the present findings refine our understanding of decision-making in psychosis, linking it to broader literature on perceptual decision processes. Our empirical results are mostly in line with recent suggestions of overweighted or hyper-precise perceptual endogenous priors in psychosis ^4,64,69,111^. Unlike some earlier studies favouring an unequivocal account of strong priors (e.g., ^48^), we find priors themselves being only slightly stronger in psychosis when compared to healthy individuals. However, their relative modulation, compared to the interpretation of sensory evidence that occurs subsequently in the decision process, was stronger. The observed reduced evidence accumulation malleability points to cognitive rigidity and impaired learning from statistical contingencies ^61,109,112^. Skepticism towards external information due to higher delusional ideation could be another conceivable reason for reduced malleability but that is not supported by our post hoc honesty ratings. More evidence for patients with psychosis attributing lower weight to trial-based sensory information is provided by reduced sensory encoding times, a difference unlikely due to motor processes often slowed by antipsychotic medication ^113^. Such hasty sampling of stimuli may represent a perceptual version of the top-down jumping-to-conclusion bias that has been linked to delusions ^114,115^. Hence, our results speak to a rigid reliance on pre-established prior beliefs, which simultaneously challenges the rival hypothesis of weaker priors alongside higher sensory weighting ^8,116^.

Reconciling these partly contradicting theories of aberrant prior expectations in psychosis will require examining endogenous priors across neural hierarchies and the entire continuum of psychotic-like experiences ^56,65^. Considering these experiences as a continuum offers explanatory power, as studies would differentiate non-clinical and clinical populations without a dichotomous classification. Our dimensional analysis based on self-reported schizotypy and delusion proneness supports this perspective and also suggests modality differences in the use of probabilistic prior information.

As for the limitations of our study, the sample size of twenty patients is not unusual for these special-needs populations but limits generalisability. Also, not all participants completed both (auditory and visual) tasks, however, we accounted for the latter by using the linear mixed models framework. Not least, residual differences in task difficulty between modalities lingered despite our efforts to equate these: The visual task appeared more challenging, as indicated by lower sensitivity and higher bias. However, the study’s main result is unaffected by this potential difference between tasks. That is, the patients’ lack of a manipulation of the evidence accumulation bias with prior information in the HDDM is independent of modality. These factors should be considered when interpreting the findings.

In conclusion, our findings advance the mechanistic understanding of how probabilistic prior information shapes perceptual decision-making across sensory modalities in both healthy individuals and those with psychosis. While healthy decision-making relies on a flexible, multi-stage process integrating changing exogenous prior information with sensory evidence, psychosis is marked by a relative reliance on pre-established prior beliefs and diminished sensory gain—giving relatively more weight to a priori expectations. Lastly, our results underscore the need for interviews about positive symptomatology in clinical populations, instead of relying on self-report scales for measuring aberrant perception.

The identified sensory gain disruption may serve as a computational marker for psychosis development and progression, with potential relevance for other disorders involving perceptual distortions. Future research should leverage neurally constrained HDDMs to disentangle the neural mechanisms underlying prior expectations in audio-visual decision-making across the psychosis spectrum.

## Methods

### Participants

We here present data from three samples: A lab-based sample of healthy control participants; a lab-based sample of participants with a diagnosis of psychosis; and a healthy online-based sample. First, we collected data from 37 individuals without a clinical diagnosis of psychosis in the laboratory of whom we included 36 participants in the final analyses (female = 25; *M*_age_ = 25.06 years, *SD*_age_ = 4.71, range = 18-37; *M*_education_ = 12.39 years, *SD*_education_ = 0.84; Table S1). One participant was excluded for repetitive behaviour. Participants were recruited using the in-house participant database ORSEE ^117^.

Second, we collected data from 20 in-patients with a diagnosis of a psychosis (F2x.x; ICD-10; ^78^) admitted to a psychosis ward at the university hospital Schleswig-Holstein (female = 9; *M*_age_ = 33.79 years, *SD*_age_ = 8.54, range = 20-49; *M*_education_ = 11 years, *SD*_education_ = 1.82). 15 out of 20 patients received prescribed antipsychotic medication around the time of testing (Chlorpromazine equivalents: *Mdn* = 388 mg, *SD* = 310 mg). Equivalents were calculated using ^118^, with non-mentioned medication converted using the daily dose method of similar active substances (e.g., Aripiprazole as substitute for Reagila)^119^. On average, the summed anticholinergic burden (ACB) of all prescribed medication score is 2.75, indicating a burden (*SD* = 1.86; www.acbcalc.com; ^120^). Though, antipsychotic medication had no direct effect on any outcome parameter in our mixed-effects analyses (see, Tables S2a-8a, 9, 10, 11a).

Lastly, we also collected and analysed data from 192 individuals without a clinical diagnosis of psychosis who took part in both tasks online via the platforms Prolific and Labvanced (female = 89; *M*_age_ = 29.26 years, *SD*_age_ = 7.62, range = 18-49; *M*_education_ = 12.34 years, *SD*_education_ = 1.24; www.prolific.com; ^121^). All participants provided written informed consent at the start and were compensated for their time financially (10€/hour) or received course credit. They self-reported of having normal or corrected to normal vision, no hearing impairment, and no neurological disorders. Participants without psychosis confirmed not having been given a diagnosis of a psychiatric illness or the presence of such a diagnosis in their first-degree relatives, and/or regular consumption of amphetamines or cannabis. The proportion of participants having learned to play a musical instrument at some point in their life is higher in individuals without psychosis (no psychosis = 78%, psychosis = 32%). This study was approved by the ethics board of the University of Lübeck in accordance with the declaration of Helsinki (certificate #21-265).

### Instruments

Before the experimental tasks, we obtained self-reported scores on participants’ proneness to schizotypal personality, hallucinations, and delusions using the Schizotypal Personality Questionnaire-Brief (SPQ-B; ^76^), Launay-Slade Hallucination Scale (LSHS-R; ^75^), and the 21-item Peter’s Delusion Inventory (PDI; ^75,77^). Additionally, participants filled in a personality inventory, which was not included in any analysis (“Big 5”; ^122^). Subsequently to the two experimental tasks, participants rated their belief in the truthfulness of the information on target probabilities provided at the start of each experimental block on a custom 5-point Likert scale (range: dishonest to honest) for each task separately. All questionnaires were administered online in their validated German form by using the experimental platform Labvanced ^121^.

### Stimuli

We generated a variety of sounds and images for two separate analogue perceptual detection tasks. Following the argument of strong parametric stimulus control for cognitive and psychophysics/acoustics experiments, all stimuli were artificially synthesised and distorted ^123^. Sounds were sampled in the form of short sound snippets (1 s in duration) from a variety of categories used in previous auditory studies ^59,124,125^ and published in official databases, such as the Jena Speaker Set (JESS) and GRID ^126,127^. These categories included human speech and voice sounds (e.g., vowels and brief sentences) as well as a wide variety of other regularly encountered sounds originating from animals, tools, and nature. The previously used stimulus set was complemented with additional speech sounds to equate numbers between stimulus categories (i.e., voice vs non-voice). If sounds obtained from the two databases JESS and GRID were longer than one second, we cut out an exemplary snippet of one second using the PRAAT software ^128^. Subsequently, we applied root-mean square (RMS) normalisation in MATLAB (version 2021a, The MathWorks, 2021, Natick, Massachusetts).

The normalised sounds were then used to synthesise sound textures, 1 second in duration and stored as .wav file with a sampling frequency of 20 kHz and a 16-bit resolution. All sounds were matched in their sound energy by manipulating their energy spectrum using RMS normalisation. Sound textures are based on an iterative approximation of selected average statistics of the temporal and spectral variation of the original sounds. The algorithm iteratively synthesises the final texture from pink noise using the sound texture toolbox in MATLAB ^73^. In the synthesis procedure, we included statistics, such as the marginal moments mean, skew, and kurtosis. We also used two sets of filters. First, a bank of 30 cochlear filters split the signal into frequency compositions with equally-spaced centre frequencies on an ERB scale ranging from 52 to 8844 Hz, and second, another set of 20 filters to measure the modulation power with equally-spaced centre frequencies on a log-scale spanning 0.5-200 Hz. To make each sound more ambiguous and discrimination more difficult, we did not include cochlear correlations that reflect distinct aspects of coordination between envelopes of different channels ^73^. We used a maximum of 45 iterations to synthesise the morphed sound textures. To achieve a parametric modulation of sound features crucial for voice perception, analogue to phase coherence modulation of images, we applied a weighting (mixing ratio: original sound/noise) to the statistics of the original sound and pink noise snippet (i.e., high noise condition: 50/50% and low noise condition: 95/5%). An online pre-test was used to determine weights resulting in sounds around the psychophysical threshold that also lead to frequent misattribution. As a result, every synthesised sound is a model-based approximation of the original sound’s structure.

Images of faces and visual wavelet textures, the analogue to sound textures, served as stimuli in the visual experiment. Colourful 3D-generated faces on black background, either 0 or 30 degrees tilted to the left or right, were derived from the MPI face database (Max Planck Institute for Biological Cybernetics, Tübingen, Germany) ^129^ and pasted on standard grey background (RGB [140 140 140]) in Photoshop. Subsequently, we grey-scaled these in MATLAB using the *rgb2gray* function (version 2021a, The MathWorks, 2021, Natick, Massachusetts). Faces were manually placed either in the centre, left or right horizontal thirds of the image. Synthesised wavelet textures served as the non-face category. These wavelet textures were iteratively generated using the visual texture toolbox in MATLAB ^130^, the visual equivalent of the sound texture toolbox used for generating sound textures (http://www.cns.nyu.edu/∼lcv/texture/). Previously, another group used the same visual toolbox with parameters set to four orientations, four scales, and a 9 x 9 spatial neighbourhood ^131^. Here, we reused these parameters. Our synthesised wavelet images had the same average statistics (i.e., contrast, kurtosis, and skew) as the used face images. The mean amplitude spectrum was then equated across images by combining the average spectrum of all images in the set. Subsequently, the images’ average luminance was equated. Lastly, we parametrically manipulated the phase coherence of both face images and wavelet textures ^132^ (custom MATLAB scripts are available from the study’s OSF repository (https://osf.io/mhqjr/). Here, we selected 17.5% (high noise), 22.5% (low noise), and 85% (catch trials) phase coherence as difficulty levels spanning the psychophysical threshold, as determined in an online pilot study (for stimulus examples, see Fig. 1C, D).

### Experimental environment

The experiment was controlled by a desktop computer (Windows 7) running Psychtoolbox-3 in MATLAB (version 2017b, The MathWorks, 2017, Natick, Massachusetts) and an external RME Fireface UC sound card. Sound was delivered binaurally via in-ear headphones (EARTONE 3A, 3M). Visual stimuli were displayed on a LED computer monitor (60Hz refresh rate; catalog #TD2421, ViewSonic). Participants were seated on a comfortable chair in a sound-attenuated booth where they kept a distance of 70 cm to the display by placing their head on a chin rest. Responses were indicated by pressing either the right or left arrow key with their right hand for the perceptual decision and subsequently one of the 1, 2, 3 or 4 keys with their left hand to indicate their confidence on a standard keyboard.

### Experimental design

Participants were first asked to complete demographic and self-report questionnaires lasting approximately 10 min. Subsequently, participants continued with the first of the counterbalanced experimental tasks. This study featured two identical detection tasks of unfamiliar speech sounds or face images in noise. Specifically, participants were asked to indicate whether a sound was generated by a human vocal tract (i.e., voice; auditory experiment) or whether an image contained a human face (visual experiment). Experimental trials, stimuli, and response button presses were identical between environments (i.e., laboratory and online). While the group without psychosis in the lab participated in both tasks on the same day, patients with psychosis were allowed to participate in each task on separate days, accounting for potentially shorter attention spans. To avoid high dropout rates online, participants could select to take part in the second experiment after a break. Each experimental task (i.e., auditory and visual) lasted approximately 30 min. The entire experiment lasted approximately 75-90 min.

At the start of each block, participants viewed an information screen that provided a truthful, probabilistic cue about the proportion of target stimuli (human voice or face) in the upcoming block (Fig. 1B). This cue was either informative by indicating fewer (P^−^; 33%) or more than half of the trials (P^+^; 66%) containing a target or entirely uninformative by indicating half of the trials containing a target (P^=^; Fig. 1B). The order of the cues was randomised, however, the cue of the first block was always uninformative. Subsequent trials followed the identical structure in both tasks irrespective of the stimulated modality and environment (Fig. 1A). First, participants viewed a blank grey screen without any content whose presentation duration was jittered between 1.5 and 2.5 seconds (inter-trial interval). Second, depending on the experiment, participants were presented with either a sound or an image for a maximum of 750 ms or until a decision was indicated by pressing one of two buttons on a standard keyboard (right and left arrow keys). In the audio-visual modalities, equating conditions comes with the challenge of stimulus dynamics unfolding differently in time. While sounds unfold over time, images are static but can be sampled dynamically over time. To allow for visual sampling, we chose a rather long image presentation time for perceptual decision-making instead of flashing the images. To accommodate potentially slower responses by patients, the response deadline for the detection decision was set to 3 seconds. If a valid decision was provided within this deadline, a one-second delay (blank grey screen) was followed by a 4-point confidence rating whose response deadline was set to 4 seconds. Participants completed nine randomised blocks consisting of 30 experimental and two catch trials per task, accumulating to 576 trials in total (i.e., per task: 135 (non-)target trials, 90 trials per cue). Catch trials served as attention checks and motivation enhancers.

Following the perceptual tasks, participants were asked to assess the truthfulness of the probabilistic cues and guess the number of difficulty levels they had encountered throughout the experiment.

### Analyses

Our data cleaning procedure required all participants, irrespective of the environment, to achieve an accuracy of at least 75% on catch trials. In the online data, according to recent guidelines for behavioural online studies ^133^, we also applied the following criteria: we only included data of participants who (i) completed both tasks (except patients), (ii) exhibited an accuracy of at least 50% on average on each task, and (iii) provided at least 80% valid trials. Valid trials are all those with a recorded button press and a confidence rating before the respective deadlines as well as a perceptual decision not faster than 250 ms. This lower bound was motivated by neuroscientific research showing 100 ms post-stimulus onset to be the lower bound of face and visual texture processing ^134^ and ∼200 ms for speech sounds ^135^. Across tasks, we removed invalid decision (lab = 0.02% of all trials, online = 0.008%), fast decision (lab = 0.002%, online = 0.0004%), and invalid confidence trials (lab = 0%, online = 0.001%). Since we had more experimental control in the lab, we retained one participant with and two without psychosis who showed an accuracy slightly below 50% in the visual task, and one patient who provided only 68% valid trials. No participant across environments failed the catch trials screening demonstrating that they understood the task. From the 20 patients with psychosis 19 took part in the visual and 17 in the auditory experiment.

For behavioural data analysis, we calculated target choice probabilities as the proportion of target choices given the number of valid trials for each prior information cue. The reference target probability varied with cue between 33%, 50%, and 66%, respectively.

Sensitivity and criterion from signal detection theory (SDT; ^71^) were calculated using custom scripts in MATLAB (version 2023b, The MathWorks, 2023, Natick, Massachusetts). This well-established theory conceptualises the outcome of a perceptual decision as a mixture of perceptual sensitivity (internal signal-to-noise ratio; SNR) and a threshold representing the internal decision criterion (bias) for the decision options. The latter is independent from the SNR as it is conceptualised as an a priori threshold before the stimulus was encountered. We calculated sensitivity (d’) and criterion (c) as follows:

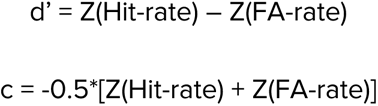

where hit rate was calculated as the proportion of correct target detections of all valid target-present trials for each cue separately. False alarm (FA) rate was calculated as the proportion of target choices on valid target-absent trials.

To quantify differences in participants’ decision behaviour and performance as well as potential links to individual traits, while accounting for inter-participant variability, we used several (generalised) linear mixed-effects models ([G]LMM; *lme4* package and *[G]LMER* functions in RStudio; ^72,136^). Specifically, we analysed the dependent variables choice probability, sensitivity, criterion, RT, hit rate, and FA rate as a function of the factors “prior information cue” (P^−^, P^=^, P^+^), “modality/task” (auditory vs visual), “stimulus noise level” (high vs low), “psychosis diagnosis” (control vs patient; only available in lab data) and antipsychotic medication (Chlorpromazine equivalent daily dose in 100mg; only available in lab data ; Tables S2-S6, S11). Additional LMMs included either individual traits (Tables S7-S10) or the sum of psychosis symptoms on the PANSS^86^, separated in positive and negative symptoms, as rated by an interviewer. The factor prior information cue was contrast coded with P^=^ as reference level. The models included participant-specific random slopes and intercepts and, where possible, stimulus-specific random intercepts. To support model convergence, where necessary, we excluded correlations between random effects and omitted random slopes. All models used the “bobyca” optimiser. To account for testing of multiple hypotheses in a given model, we applied an FDR-correction to all reported p-values, as implemented in the tab_model function of the *sjPlot* package (version 2.8.15; ^137^).

Results of choice probability, hit and FA rate were based on single-trial information and logistic GLMMs. The model quantifying continuous RT data also used single-trial information but analysed effects on log(RT) in an LMM with restricted maximum likelihood estimation (REML). All models quantifying the SDT metrics sensitivity and criterion are based on one value per participant and experimental condition of interest. The results were obtained using separate LMMs with REML (*LMER* function; ^72,136^).

To evaluate participants’ trust in the truthfulness of our prior information cues, we used cumulative link mixed models (using the *clmm* function from the *ordinal* package; version 2023.12-4.1, ^138^), which allow for estimating ordinal data. Similar to our (G)LMMs, we included a participant-specific random intercept and used the default logit link function.

Differences in individual traits (i.e., proneness to schizotypy, hallucinations, and delusions) between groups (participants with vs without psychosis) and the testing environments (laboratory vs online) were quantified in RStudio using negative binomial generalised linear models (*glm.nb* function of the *MASS* package; ^139^). We used contrast coding with the group of participants without psychosis who participated in the lab as the reference.

We fit Bayesian hierarchical drift diffusion models (HDDMs) to RT distributions as a function of choice (i.e., stimulus coding) using a docker image of the HDDM python toolbox (^57,58,90^; Fig. 4C, D). Specifically, we based this model on findings from previous comparisons of analogue auditory and visual perceptual decisions and ambiguous image detection in psychosis-prone non-clinical samples ^45,89^. Leveraging these models allowed us to differentiate between different constituents of the decision-making process, with a specific focus on bias parameters as a function of prior information and sensory modality. To benefit from hierarchical dependencies, we fit two separate but identical models, one to participants without (N=36) and another one to those with psychosis (N=20).

We included all standard parameters of the HDDM but omitted inter-trial-variation parameters, as the variation between blocks of trials, and not between trials, was most pertinent to our question. As well, these parameters often provide limited gain, while negatively impacting convergence. Hence, we evaluated sensory encoding and response execution (nondecision time, *nDT*), quality of the evidence accumulation (drift rate, *v*), amount of evidence required for a choice or general response caution (boundary separation, *a*), internal bias towards one of the two choice options without considering stimulus evidence (starting point, *z*), and a bias of the evidence accumulation process (drift criterion, dc). Particularly, the latter two parameters are complementary and allow the hierarchical drift-diffusion model to capture nuanced aspects of a priori response and sensory evidence processing biases in human decision-making. To account for specific variations of prior information (P^−^, P^=^, P^+^), task modality (auditory vs visual), and stimulus noise (low vs high), in line with our SDT results, we included dependencies of several parameters. That is, *nDT* depended on the stimulated modality due to different temporal stimulus properties, *z* and *dc* depended on prior information and modality, and *v* on an interaction of prior information and stimulus noise. The proportion of RT outliers (*p_outlier*) was empirically fixed to 1.5%, based on estimations of this proportion as a free parameter. The *a* parameter indicates general differences between speed vs. accuracy response strategies set at the start of a task. It most sensibly varies between participants with and without psychosis but not the experimental conditions. Therefore, *a* was allowed to vary freely.

More specifically, we used the HDDM.StimCoding function to estimate drift rates in relation to the presented stimulus (i.e., target-present vs target-absent). Sampling parameters were set to three chains with 6000 samples, 2000 samples burn-in, a thinning factor of 4 samples, and three chains resulting in evidence based on 1000 final samples per chain. Models were diagnosed for convergence visually as well as using the Gelman-Rubin (R^ ; threshold <= 1.01; nDT for one control = 1.0119, all other parameters < 1.008; patient: all parameters < 1.006) and effective sample size statistics (ESS; threshold >= 400; control: all parameters > 506, patients: one subject nDT = 285 but R^ < 1.01). Subsequently, we used the DIC ^140^ to select the best fitting converging model, which resulted in the model including dependent *v*, *z* and *dc* parameters. Models excluding the dependency of the drift criterion on prior information and modality fared worse than those excluding a starting point depending on the same conditions (control: DIC_full_ = 31000, DIC_fixed-z_ = 31212, DIC_fixed-dc_ = 31311; patient: DIC_full_ = 18278, DIC_fixed-z_ = 18311, DIC_fixed-dc_ = 18436; Tables S14). Lastly, we tested for the probability of a statistically-relevant difference by comparing the 95% highest density intervals (HDIs) of relevant posterior distributions and computed the probability of direction that one posterior distribution is larger/smaller than the mean of another.

## Acknowledgements

We would like to thank Malte Wöstmann for his comments on experimental design and earlier versions of the manuscript. During the preparation of this work the authors used the Generative AI service ChatGPT, v. 4o, as provided by OpenAI Inc. for light editing of certain paragraphs. After using this tool, the authors reviewed and edited the content as needed and take full responsibility for the content of the publication.

## Competing interests

none

## Funding information

European Research Council (ERC-CoG-2014 No. 646696 “AUDADAPT”) awarded to JO (2016– 2021)

## CREDIT author contributions

Conceptualisation: L.F., S.E., C.A., J.E., S.T., and J.O.

Data curation: L.F., S.E. and H.S.

Formal analysis: L.F., I.D., J.K., N.K., and S.T.

Funding acquisition: C.A., S.B., and J.O.

Investigation: L.F., S.E., C.L., R.L., and H.S.

Methodology: L.F., I.D., J.K., N.K. and S.T.

Project administration: L.F., C.A., and J.O.

Resources: L.F., C.A., J.E., S.B., and J.O.

Software: L.F., L.M.S., and S.T.

Supervision: C.A., S.B., and J.O.

Validation: L.F.

Visualisation: L.F. and J.O.

Writing - original draft: L.F., H.S., and J.O.

Writing - review & editing: L.F., S.E., C.A., I.D., J.E., J.K., C.L., R.L., L.M.S., H.S., N.K., S.T., S.B., and J.O.

## Data availability

Data and code will be available from the project’s Open Science Framework repository https://osf.io/mhqjr/.

## Notes

### Competing Interest Statement

The authors have declared no competing interest.

## References

1. Knill, D. C. & Pouget, A. The Bayesian brain: the role of uncertainty in neural coding and computation. Trends Neurosci. 27, 712–719 (2004).

2. Kersten, D., Mamassian, P. & Yuille, A. Object Perception as Bayesian Inference. Annu. Rev. Psychol. 55, 271–304 (2004).

3. McCutcheon, R., Marques, T. & Howes, O. Schizophrenia—An Overview. JAMA Psychiatry 77, 201–210 (2020).

4. Powers, A. R. et al. A Computational Account of the Development and Evolution of Psychotic Symptoms. Biol. Psychiatry 97, 117–127 (2025).

5. Horga, G. & Abi-Dargham, A. An integrative framework for perceptual disturbances in psychosis. Nat. Rev. Neurosci. 20, 763–778 (2019).

6. Fletcher, P. C. & Frith, C. D. Perceiving is believing: a Bayesian approach to explaining the positive symptoms of schizophrenia. Nat. Rev. Neurosci. 10, 48–58 (2009).

7. Adams, R. A., Stephan, K. E., Brown, H. R., Frith, C. D. & Friston, K. J. The Computational Anatomy of Psychosis. Front. Psychiatry 4, (2013).

8. Sterzer, P. et al. The Predictive Coding Account of Psychosis. Biol. Psychiatry 84, 634–643 (2018).

9. Fromm, S. P. et al. Neural correlates of uncertainty processing in psychosis spectrum disorder. Brain Commun. 7, fcaf073 (2025).

10. Gold, J. I. & Shadlen, M. N. Neural computations that underlie decisions about sensory stimuli. Trends Cogn. Sci. 5, 10–16 (2001).

11. Heekeren, H. R., Marrett, S. & Ungerleider, L. G. The neural systems that mediate human perceptual decision making. Nat. Rev. Neurosci. 9, 467–479 (2008).

12. Yuille, A. & Kersten, D. Vision as Bayesian inference: analysis by synthesis? Trends Cogn. Sci. 10, 301–308 (2006).

13. Summerfield, C. & De Lange, F. P. Expectation in perceptual decision making: neural and computational mechanisms. Nat. Rev. Neurosci. 15, 745–756 (2014).

14. Summerfield, C. & Egner, T. Expectation (and attention) in visual cognition. Trends Cogn. Sci. 13, 403–409 (2009).

15. Bévalot, C. & Meyniel, F. A dissociation between the use of implicit and explicit priors in perceptual inference. Commun. Psychol. 2, 111 (2024).

16. Bogacz, R., Brown, E., Moehlis, J., Holmes, P. & Cohen, J. D. The physics of optimal decision making: A formal analysis of models of performance in two-alternative forced-choice tasks. Psychol. Rev. 113, 700–765 (2006).

17. Oliva, A. & Torralba, A. The role of context in object recognition. Trends Cogn. Sci. 11, 520– 527 (2007).

18. Mulder, M. J., Wagenmakers, E.-J., Ratcliff, R., Boekel, W. & Forstmann, B. U. Bias in the Brain: A Diffusion Model Analysis of Prior Probability and Potential Payoff. J. Neurosci. 32, 2335–2343 (2012).

19. Weilnhammer, V. A., Stuke, H., Sterzer, P. & Schmack, K. The Neural Correlates of Hierarchical Predictions for Perceptual Decisions. J. Neurosci. 38, 5008–5021 (2018).

20. White, C. N. & Poldrack, R. A. Decomposing bias in different types of simple decisions. J. Exp. Psychol. Learn. Mem. Cogn. 40, 385–398 (2014).

21. Kok, P., Mostert, P. & De Lange, F. P. Prior expectations induce prestimulus sensory templates. Proc. Natl. Acad. Sci. 114, 10473–10478 (2017).

22. Puri, A. M., Wojciulik, E. & Ranganath, C. Category expectation modulates baseline and stimulus-evoked activity in human inferotemporal cortex. Brain Res. 1301, 89–99 (2009).

23. Wyart, V., Nobre, A. C. & Summerfield, C. Dissociable prior influences of signal probability and relevance on visual contrast sensitivity. Proc. Natl. Acad. Sci. 109, 3593–3598 (2012).

24. Kelly, S. P., Corbett, E. A. & O’Connell, R. G. Neurocomputational mechanisms of prior-informed perceptual decision-making in humans. *Nat*. Hum. Behav. 5, 467–481 (2021).

25. Wiech, K. et al. Influence of prior information on pain involves biased perceptual decision-making. Curr. Biol. 24, R679–R681 (2014).

26. Ratcliff, R., Smith, P. L., Brown, S. D. & McKoon, G. Diffusion Decision Model: Current Issues and History. Trends Cogn. Sci. 20, 260–281 (2016).

27. Cerracchio, E., Miletić, S. & Forstmann, B. U. Modelling decision-making biases. Front. Comput. Neurosci. 17, 1222924 (2023).

28. Cravo, A. M., Rohenkohl, G., Wyart, V. & Nobre, A. C. Temporal Expectation Enhances Contrast Sensitivity by Phase Entrainment of Low-Frequency Oscillations in Visual Cortex. J. Neurosci. 33, 4002–4010 (2013).

29. De Gee, J. W. et al. Dynamic modulation of decision biases by brainstem arousal systems. eLife 6, e23232 (2017).

30. Diaz, J. A., Pisauro, M. A., Delis, I. & Philiastides, M. G. Prior probability biases perceptual choices by modulating the accumulation rate, rather than the baseline, of decision evidence. Imaging Neurosci. 2, 1–19 (2024).

31. Hanks, T. D., Mazurek, M. E., Kiani, R., Hopp, E. & Shadlen, M. N. Elapsed Decision Time Affects the Weighting of Prior Probability in a Perceptual Decision Task. J. Neurosci. 31, 6339–6352 (2011).

32. Kloosterman, N. A. et al. Humans strategically shift decision bias by flexibly adjusting sensory evidence accumulation. eLife 8, e37321 (2019).

33. Rungratsameetaweemana, N., Itthipuripat, S., Salazar, A. & Serences, J. T. Expectations Do Not Alter Early Sensory Processing during Perceptual Decision-Making. J. Neurosci. 38, 5632–5648 (2018).

34. Walsh, K., McGovern, D. P., Dully, J., Kelly, S. & O’Connell, R. Prior probability cues bias sensory encoding with increasing task exposure. Preprint at 10.7554/eLife.91135.2 (2024).

35. Dunovan, K. E., Tremel, J. J. & Wheeler, M. E. Prior probability and feature predictability interactively bias perceptual decisions. Neuropsychologia 61, 210–221 (2014).

36. Dunovan, K. E. & Wheeler, M. E. Computational and neural signatures of pre and post-sensory expectation bias in inferior temporal cortex. Sci. Rep. 8, 13256 (2018).

37. Retzler, C., Boehm, U., Cai, J., Cochrane, A. & Manning, C. Prior information use and response caution in perceptual decision-making: No evidence for a relationship with autistic-like traits. Q. J. Exp. Psychol. 74, 1953–1965 (2021).

38. van Ravenzwaaij, D., Mulder, M. J., Tuerlinckx, F. & Wagenmakers, E.-J. Do the dynamics of prior information depend on task context? An analysis of optimal performance and an empirical test. CrossRef Listing Deleted DOIs 3, 132 (2012).

39. O’Connell, R. G. & Kelly, S. P. Neurophysiology of Human Perceptual Decision-Making. Annu. Rev. Neurosci. 44, 495–516 (2021).

40. Levine, S. M. & Schwarzbach, J. V. Cross-decoding supramodal information in the human brain. Brain Struct. Funct. 223, 4087–4098 (2018).

41. O’Connell, R. G., Dockree, P. M. & Kelly, S. P. A supramodal accumulation-to-bound signal that determines perceptual decisions in humans. Nat. Neurosci. 15, 1729–1735 (2012).

42. Spitzer, B., Blankenburg, F. & Summerfield, C. Rhythmic gain control during supramodal integration of approximate number. NeuroImage 129, 470–479 (2016).

43. Brunton, B. W., Botvinick, M. M. & Brody, C. D. Rats and Humans Can Optimally Accumulate Evidence for Decision-Making. Science 340, 95–98 (2013).

44. Hupé, J.-M., Joffo, L.-M. & Pressnitzer, D. Bistability for audiovisual stimuli: Perceptual decision is modality specific. J. Vis. 8, 1 (2008).

45. Mulder, M. J. et al. The speed and accuracy of perceptual decisions in a random-tone pitch task. Atten. Percept. Psychophys. 75, 1048–1058 (2013).

46. Neri, P. How inherently noisy is human sensory processing? Psychon. Bull. Rev. 17, 802–808 (2010).

47. Tamber-Rosenau, B. J., Dux, P. E., Tombu, M. N., Asplund, C. L. & Marois, R. Amodal Processing in Human Prefrontal Cortex. J. Neurosci. 33, 11573–11587 (2013).

48. Teufel, C. et al. Shift toward prior knowledge confers a perceptual advantage in early psychosis and psychosis-prone healthy individuals. Proc. Natl. Acad. Sci. 112, 13401–13406 (2015).

49. Alderson-Day, B. et al. Susceptibility to auditory hallucinations is associated with spontaneous but not directed modulation of top-down expectations for speech. Neurosci. Conscious. 2022, niac002 (2022).

50. Silverstein, S. M. & Lai, A. The Phenomenology and Neurobiology of Visual Distortions and Hallucinations in Schizophrenia: An Update. Front. Psychiatry 12, (2021).

51. Morgan, M. J. & Ward, R. Conditions for motion flow in dynamic visual noise. Vision Res. 20, 431–435 (1980).

52. Mathias, S. R. et al. The Processing-Speed Impairment in Psychosis Is More Than Just Accelerated Aging. Schizophr. Bull. sbw168 (2017) doi:10.1093/schbul/sbw168.

53. Shen, C., Calvin, O. L., Rawls, E., Redish, A. D. & Sponheim, S. R. Clarifying Cognitive Control Deficits in Psychosis via Drift Diffusion Modeling and Attractor Dynamics. Schizophr. Bull. 50, 1357–1370 (2024).

54. Klatt, L.-I. et al. Unraveling the Relation between EEG Correlates of Attentional Orienting and Sound Localization Performance: A Diffusion Model Approach. J. Cogn. Neurosci. 32, 945– 962 (2020).

55. Liu, A. S. K., Tsunada, J., Gold, J. I. & Cohen, Y. E. Temporal Integration of Auditory Information Is Invariant to Temporal Grouping Cues. eneuro 2, ENEURO.0077-14.2015 (2015).

56. Heinz, A. et al. Towards a Unifying Cognitive, Neurophysiological, and Computational Neuroscience Account of Schizophrenia. Schizophr. Bull. 45, 1092–1100 (2019).

57. Pan, W. et al. A Hitchhiker’s Guide to Bayesian Hierarchical Drift-Diffusion Modeling with dockerHDDM. Preprint at (2024).

58. Wiecki, T. V., Sofer, I. & Frank, M. J. HDDM: Hierarchical Bayesian estimation of the Drift-Diffusion Model in Python. *Front*. Neuroinformatics 7, (2013).

59. Erb, J., Kreitewolf, J., Pinheiro, A. P. & Obleser, J. Aberrant Perceptual Judgments on Speech-Relevant Acoustic Features in Hallucination-Prone Individuals. Schizophr. Bull. Open 1, sgaa059 (2020).

60. Tarasi, L., Martelli, M. E., Bortoletto, M., Di Pellegrino, G. & Romei, V. Neural Signatures of Predictive Strategies Track Individuals Along the Autism-Schizophrenia Continuum. Schizophr. Bull. 49, 1294–1304 (2023).

61. Eckert, A.-L., Gounitski, Y., Guggenmos, M. & Sterzer, P. Cross-Modality Evidence for Reduced Choice History Biases in Psychosis-Prone Individuals. Schizophr. Bull. 49, 397– 406 (2023).

62. Abrahamyan, A., Silva, L. L., Dakin, S. C., Carandini, M. & Gardner, J. L. Adaptable history biases in human perceptual decisions. Proc. Natl. Acad. Sci. 113, (2016).

63. Linszen, M. M. J. et al. Occurrence and phenomenology of hallucinations in the general population: A large online survey. Schizophrenia 8, 41 (2022).

64. Corlett, P. R. et al. Hallucinations and Strong Priors. Trends Cogn. Sci. 23, 114–127 (2019).

65. Haarsma, J. et al. Influence of prior beliefs on perception in early psychosis: Effects of illness stage and hierarchical level of belief. J. Abnorm. Psychol. 129, 581–598 (2020).

66. Haarsma, J., Kok, P. & Browning, M. The promise of layer-specific neuroimaging for testing predictive coding theories of psychosis. Schizophr. Res. 245, 68–76 (2022).

67. Keller, G. B. & Sterzer, P. Predictive Processing: A Circuit Approach to Psychosis. Annu. Rev. Neurosci. 47, 85–101 (2024).

68. Powers, A. R., Kelley, M. & Corlett, P. R. Hallucinations as Top-Down Effects on Perception. Biol. Psychiatry Cogn. Neurosci. Neuroimaging 1, 393–400 (2016).

69. Powers, A. R., Mathys, C. & Corlett, P. R. Pavlovian conditioning–induced hallucinations result from overweighting of perceptual priors. Science 357, 596–600 (2017).

70. Stuke, H., Weilnhammer, V. A., Sterzer, P. & Schmack, K. Delusion Proneness is Linked to a Reduced Usage of Prior Beliefs in Perceptual Decisions. Schizophr. Bull. (2018) doi:10.1093/schbul/sbx189.

71. Green, D. M. & Swets, J. A. Signal Detection Theory and Psychophysics. xiii, 479 (Robert E. Krieger, Oxford, England, 1974).

72. Baayen, R. H., Davidson, D. J. & Bates, D. M. Mixed-effects modeling with crossed random effects for subjects and items. J. Mem. Lang. 59, 390–412 (2008).

73. McDermott, J. H. & Simoncelli, E. P. Sound Texture Perception via Statistics of the Auditory Periphery: Evidence from Sound Synthesis. Neuron 71, 926–940 (2011).

74. Allen, M., Poggiali, D., Whitaker, K., Marshall, T. R. & Kievit, R. A. Raincloud plots: a multi-platform tool for robust data visualization. Wellcome Open Res. 4, 63 (2019).

75. Lincoln, T. M., Keller, E. & Rief, W. Die Erfassung von Wahn und Halluzinationen in der Normalbevölkerung: Deutsche Adaptationen des Peters et al. Delusions Inventory (PDI) und der Launay Slade Hallucination Scale (LSHS-R). Diagnostica 55, 29–40 (2009).

76. Raine, A. & Benishay, D. The SPQ-B: A Brief Screening Instrument for Schizotypal Personality Disorder. J. Personal. Disord. 9, 346–355 (1995).

77. Peters, E., Joseph, S., Day, S. & Garety, P. Measuring Delusional Ideation: The 21-Item Peters et al. Delusions Inventory (PDI). Schizophr. Bull. 30, 1005–1022 (2004).

78. World Health Organization. ICD-10: international statistical classification of diseases and related health problems: tenth revision. (World Health Organization, Geneva, 2004).

79. Crow, T. J. The Continuum of Psychosis and its Implication for the Structure of the Gene. Br. J. Psychiatry 149, 419–429 (1986).

80. Strauss, J. S. Hallucinations and Delusions as Points on Continua Function: Rating Scale Evidence. Arch. Gen. Psychiatry 21, 581 (1969).

81. Van Os, J., Linscott, R. J., Myin-Germeys, I., Delespaul, P. & Krabbendam, L. A systematic review and meta-analysis of the psychosis continuum: evidence for a psychosis proneness– persistence–impairment model of psychotic disorder. Psychol. Med. 39, 179–195 (2009).

82. Van Os, J. & Reininghaus, U. Psychosis as a transdiagnostic and extended phenotype in the general population. World Psychiatry 15, 118–124 (2016).

83. Swets, J. A., Tanner, W. P. & Birdsall, A. G. Decision processes in perception. 68, 301–340 (1961).

84. Terman, M. & Terman, J. S. Concurrent variation of response bias and sensitivity in an operant-psychophysical test. Percept. Psychophys. 11, 428–432 (1972).

85. DeRosse, P. & Karlsgodt, K. H. Examining the Psychosis Continuum. Curr. Behav. Neurosci. Rep. 2, 80–89 (2015).

86. Kay, S. R., Fiszbein, A. & Opler, L. A. The Positive and Negative Syndrome Scale (PANSS) for Schizophrenia. Schizophr. Bull. 13, 261–276 (1987).

87. Fetsch, C. R., Kiani, R. & Shadlen, M. N. Predicting the Accuracy of a Decision: A Neural Mechanism of Confidence. Cold Spring Harb. Symp. Quant. Biol. 79, 185–197 (2014).

88. Limongi, R., Bohaterewicz, B., Nowicka, M., Plewka, A. & Friston, K. J. Knowing when to stop: Aberrant precision and evidence accumulation in schizophrenia. Schizophr. Res. 197, 386–391 (2018).

89. Davies, D. J., Teufel, C. & Fletcher, P. C. Anomalous Perceptions and Beliefs Are Associated With Shifts Toward Different Types of Prior Knowledge in Perceptual Inference. Schizophr. Bull. 44, 1245–1253 (2018).

90. Chuan-Peng, H., Geng, H., Zhang, L., Fengler, A. & Frank, M. J. A hitchhiker’s guide to bayesian hierarchical drift-diffusion modeling with dockerHDDM. Preprint at (2022).

91. Roitman, J. D. & Shadlen, M. N. Response of Neurons in the Lateral Intraparietal Area during a Combined Visual Discrimination Reaction Time Task. J. Neurosci. 22, 9475–9489 (2002).

92. Kloosterman, N. A., Kosciessa, J. Q., Lindenberger, U., Fahrenfort, J. J. & Garrett, D. D. Boosts in brain signal variability track liberal shifts in decision bias. eLife 9, e54201 (2020).

93. Rahnev, D., Lau, H. & De Lange, F. P. Prior Expectation Modulates the Interaction between Sensory and Prefrontal Regions in the Human Brain. J. Neurosci. 31, 10741–10748 (2011).

94. Tulver, K., Aru, J., Rutiku, R. & Bachmann, T. Individual differences in the effects of priors on perception: A multi-paradigm approach. Cognition 187, 167–177 (2019).

95. Weilnhammer, V., Stuke, H., Standvoss, K. & Sterzer, P. Sensory processing in humans and mice fluctuates between external and internal modes. PLOS Biol. 21, e3002410 (2023).

96. Delis, I., Ince, R. A. A., Sajda, P. & Wang, Q. Neural Encoding of Active Multi-Sensing Enhances Perceptual Decision-Making via a Synergistic Cross-Modal Interaction. J. Neurosci. 42, 2344–2355 (2022).

97. Franzen, L., Delis, I., De Sousa, G., Kayser, C. & Philiastides, M. G. Auditory information enhances post-sensory visual evidence during rapid multisensory decision-making. Nat. Commun. 11, 5440 (2020).

98. Leite, F. P. & Ratcliff, R. What cognitive processes drive response biases? A diffusion model analysis. Judgm. Decis. Mak. 6, 651–687 (2011).

99. Moran, R. Optimal decision making in heterogeneous and biased environments. Psychon. Bull. Rev. 22, 38–53 (2015).

100. Friston, K. A theory of cortical responses. Philos. Trans. R. Soc. B Biol. Sci. 360, 815–836 (2005).

101. De Lange, F. P., Rahnev, D. A., Donner, T. H. & Lau, H. Prestimulus Oscillatory Activity over Motor Cortex Reflects Perceptual Expectations. J. Neurosci. 33, 1400–1410 (2013).

102. Drugowitsch, J., Wyart, V., Devauchelle, A.-D. & Koechlin, E. Computational Precision of Mental Inference as Critical Source of Human Choice Suboptimality. Neuron 92, 1398–1411 (2016).

103. Vilares, I. & Kording, K. Bayesian models: the structure of the world, uncertainty, behavior, and the brain. Ann. N. Y. Acad. Sci. 1224, 22–39 (2011).

104. Chen, Y., Norton, D., Ongur, D. & Heckers, S. Inefficient Face Detection in Schizophrenia. Schizophr. Bull. 34, 367–374 (2007).

105. Christensen, B. K., Spencer, J. M. Y., King, J. P., Sekuler, A. B. & Bennett, P. J. Noise as a mechanism of anomalous face processing among persons with Schizophrenia. Front. Psychol. 4, (2013).

106. Lee, S.-H., Chung, Y.-C., Yang, J.-C., Kim, Y.-K. & Suh, K.-Y. Abnormal speech perception in schizophrenia with auditory hallucinations. Acta Neuropsychiatr. 16, 154–159 (2014).

107. Moseley, P. et al. Continuities and Discontinuities in the Cognitive Mechanisms Associated With Clinical and Nonclinical Auditory Verbal Hallucinations. Clin. Psychol. Sci. 10, 752–766 (2022).

108. She, S. et al. Deficits in prosodic speech-in-noise recognition in schizophrenia patients and its association with psychiatric symptoms. BMC Psychiatry 24, 864 (2024).

109. Cassidy, C. M. et al. A Perceptual Inference Mechanism for Hallucinations Linked to Striatal Dopamine. Curr. Biol. 28, 503–514.e4 (2018).

110. Benrimoh, D. et al. Evidence for Reduced Sensory Precision and Increased Reliance on Priors in Hallucination-Prone Individuals in a General Population Sample. Schizophr. Bull. 50, 349–362 (2024).

111. Schmack, K., Bosc, M., Ott, T., Sturgill, J. F. & Kepecs, A. Striatal dopamine mediates hallucination-like perception in mice. Science 372, eabf4740 (2021).

112. Sheffield, J. M., Karcher, N. R. & Barch, D. M. Cognitive Deficits in Psychotic Disorders: A Lifespan Perspective. Neuropsychol. Rev. 28, 509–533 (2018).

113. Morrens, M., Hulstijn, W. & Sabbe, B. Psychomotor Slowing in Schizophrenia. Schizophr. Bull. 33, 1038–1053 (2007).

114. Gawęda, Ł., Staszkiewicz, M. & Balzan, R. P. The relationship between cognitive biases and psychological dimensions of delusions: The importance of jumping to conclusions. J. Behav. Ther. Exp. Psychiatry 56, 51–56 (2017).

115. Sheffield, J. M., Smith, R., Suthaharan, P., Leptourgos, P. & Corlett, P. R. Relationships between cognitive biases, decision-making, and delusions. Sci. Rep. 13, 9485 (2023).

116. Weilnhammer, V. et al. Psychotic Experiences in Schizophrenia and Sensitivity to Sensory Evidence. Schizophr. Bull. 46, 927–936 (2020).

117. Greiner, B. Subject pool recruitment procedures: organizing experiments with ORSEE. J. Econ. Sci. Assoc. 1, 114–125 (2015).

118. Gardner, D. M. et al. International Consensus Study of Antipsychotic Dosing. Am J Psychiatry (2010).

119. Leucht, S., Samara, M., Heres, S. & Davis, J. M. Dose Equivalents for Antipsychotic Drugs: The DDD Method: Table 1. Schizophr. Bull. 42, S90–S94 (2016).

120. Kiesel, E. K., Hopf, Y. M. & Drey, M. An anticholinergic burden score for German prescribers: score development. BMC Geriatr. 18, 239 (2018).

121. Finger, H., Goeke, C., Diekamp, D., Standvoß, K. & König, P. LabVanced: a unified JavaScript framework for online studies. in (Cologne, 2017).

122. Hahn, E., Gottschling, J. & Spinath, F. M. Short measurements of personality – Validity and reliability of the GSOEP Big Five Inventory (BFI-S). J. Res. Personal. 46, 355–359 (2012).

123. Rust, N. C. & Movshon, J. A. In praise of artifice. Nat. Neurosci. 8, 1647–1650 (2005).

124. Norman-Haignere, S., Kanwisher, N. G. & McDermott, J. H. Distinct Cortical Pathways for Music and Speech Revealed by Hypothesis-Free Voxel Decomposition. Neuron 88, 1281– 1296 (2015).

125. Santoro, R. et al. Reconstructing the spectrotemporal modulations of real-life sounds from fMRI response patterns. Proc. Natl. Acad. Sci. 114, 4799–4804 (2017).

126. Cooke, M., Barker, J., Cunningham, S. & Shao, X. An audio-visual corpus for speech perception and automatic speech recognition. J. Acoust. Soc. Am. 120, 2421–2424 (2006).

127. Zäske, R., Skuk, V. G., Golle, J. & Schweinberger, S. R. The Jena Speaker Set (JESS)—A database of voice stimuli from unfamiliar young and old adult speakers. Behav. Res. Methods 52, 990–1007 (2020).

128. Boersma, P. & van Heuven, V. Speak and unSpeak with PRAAT. Glot International 5, (2001).

129. Troje, N. F. & Bülthoff, H. H. Face recognition under varying poses: The role of texture and shape. Vision Res. 36, 1761–1771 (1996).

130. Portilla, J. & Simoncelli, E. P. A Parametric Texture Model Based on Joint Statistics of Complex Wavelet Coefficients. Int. J. Comput. Vis. 40, 49–71 (2000).

131. Rousselet, G. A., Pernet, C. R., Bennett, P. J. & Sekuler, A. B. Parametric study of EEG sensitivity to phase noise during face processing. BMC Neurosci. 9, 98 (2008).

132. Ales, J. M., Farzin, F., Rossion, B. & Norcia, A. M. An objective method for measuring face detection thresholds using the sweep steady-state visual evoked response. J. Vis. 12, 18–18 (2012).

133. Gagné, N. & Franzen, L. How to Run Behavioural Experiments Online: Best Practice Suggestions for Cognitive Psychology and Neuroscience. *Swiss Psychol*. Open 3, 1 (2023).

134. Bieniek, M. M., Bennett, P. J., Sekuler, A. B. & Rousselet, G. A. A robust and representative lower bound on object processing speed in humans. Eur. J. Neurosci. 44, 1804–1814 (2016).

135. Schall, S., Kiebel, S. J., Maess, B. & Von Kriegstein, K. Early auditory sensory processing of voices is facilitated by visual mechanisms. NeuroImage 77, 237–245 (2013).

136. Bates, D., Mächler, M., Bolker, B. & Walker, S. Fitting Linear Mixed-Effects Models Using lme4. J. Stat. Softw. 67, (2015).

137. Lüdecke, D. sjPlot: Data Visualization for Statistics in Social Science. (2024).

138. Christensen, R. H. B. ordinal---Regression Models for Ordinal Data. (2023).

139. Venables, W. N. & Ripley, B. D. Modern Applied Statistics with S. (Springer, New York, 2002).

140. Spiegelhalter, D. J., Best, N. G., Carlin, B. P. & Van Der Linde, A. Bayesian Measures of Model Complexity and Fit. J. R. Stat. Soc. Ser. B Stat. Methodol. 64, 583–639 (2002)

