## Supplementary Materials for "Prior Information Shapes Perceptual Evidence Accumulation Dynamics Differentially in Psychosis"

**TABLE S1. Comparison of sample characteristics.** All averages of self-report questionnaire scores (i.e., SPQ, LSHS-R, PDI) are 5% trimmed means.

| **Parameter** | **Without psychosis (lab, N = 36)** | **Without psychosis (online, N = 192)** | **With psychosis (lab, N = 20)** |
|---|---|---|---|
| Gender | female = 25 | female = 89 | female = 9 |
| Age (years) | *M* = 25.06, *SD* = 4.71, range = 18-37 | *M* = 29.26, *SD* = 7.62, range = 18-49 | *M* = 33.79, *SD* = 8.54, range = 20-49 |
| Education (years) | *M* = 12.39, *SD* = 0.84 | *M* = 12.34, *SD* = 1.24 | *M* = 11, *SD* = 1.82 |
| Antipsychotic Medication (Chlorpromazine equivalents, mg) | 0 | 0 | *Mdn* = 380, *SD* = 310 |
| Trust in prior information (7-point Likert scale) | *M* = 2.43, *SD* = 1.03 | *M* = 3.08, *SD* = 1.24 | *M* = 3.09, *SD* = 1.07 |
| Schizotypy (SPQ-B) | *M* = 6.82, *SD* = 4.26, range = 0-15 | *M* = 6.91, *SD* = 4.31, range = 0-20 | *M* = 9.40, *SD* = 4.83, range = 2-18 |
| Delusions (PDI-21) | *M* = 3.68, *SD* = 2.97, range = 0-10 | *M* = 3.22, *SD* = 2.69, range = 0-12 | *M* = 7.15, *SD* = 4.99, range = 0-18 |
| Hallucinations (LSHS-R) | *M* = 14.18, *SD* = 9.15, range = 0-41 | *M* = 10.92, *SD* = 7.75, range = 0-40 | *M* = 15.05, *SD* = 11.53, range = 0-37 |

### Trait scores do not differ with previous psychological treatment

A small number of participants without a clinical diagnosis of a psychotic disorder (lab: 5/36; online: 28/192) reported having received previous psychological treatment at some point in their lives due to non-psychotic problems. Therefore, we examined non-clinical participants on potential trait differences as a function of previous treatment. We did not observe any significant difference on schizotypal personality, hallucination or delusion proneness in our lab sample (SPQ: $t(34) = 0.08$, $p = .9387$; PDI: $t(34) = 0.04$, $p = .9683$; LSHS: $t(34) = 0.12$, $p = .9031$). Hence, we considered this group of participants without a diagnosis of psychosis one healthy control group for all statistical analyses.

**TABLE S2a. Results of GLMM regression analysis predicting choice probability (lab data).**

| | Choice probability for face/voice | | |
|---|---|---|---|
| Predictors | Odds Ratios | CI | p |
| (Intercept) | 0.33 | 0.24 – 0.45 | **< .001** |
| Chlorpromazine equivalent 100mg | 1.17 | 1.02 – 1.34 | .400 |
| Prior cue (P- vs P=) | 0.64 | 0.49 – 0.84 | **.006** |
| Prior cue (P+ vs P=) | 1.34 | 1.02 – 1.76 | .075 |
| Patient | 0.46 | 0.23 – 0.93 | .075 |
| Task | 0.60 | 0.45 – 0.82 | **.006** |
| Stimulus noise | 0.52 | 0.41 – 0.65 | **< .001** |
| Prior cue (P- vs P=) × task | 1.24 | 0.72 – 2.12 | .0593 |
| Prior cue (P+ vs P=) × task | 1.40 | 0.81 – 2.39 | .385 |
| Prior cue (P- vs P=) × stimulus noise | 1.13 | 0.66 – 1.95 | .696 |
| Prior cue (P+ vs P=) × stimulus noise | 1.15 | 0.67 – 1.96 | .696 |
| Prior cue (P- vs P=) × patient | 1.13 | 0.96 – 1.33 | .314 |
| Prior cue (P+ vs P=) × patient | 0.93 | 0.79 – 1.09 | .502 |
| Patient × task | 0.75 | 0.47 – 1.21 | .385 |
| Patient × stimulus noise | 1.05 | 0.84 – 1.30 | .696 |
| Task × stimulus noise | 1.09 | 0.70 – 1.70 | .696 |
| Random Effects | | | |
| $\sigma^2$ | 3.29 | | |
| $\tau_{00\ \mathrm{stim\_ID.1}}$ | 1.57 | | |
| $\tau_{00\ \mathrm{sub\_ID.2}}$ | 0.82 | | |
| $\tau_{11\ \mathrm{stim\_ID.patient}}$ | 0.02 | | |
| $\tau_{11\ \mathrm{sub\_ID.stimulus\_noise}}$ | 0.09 | | |
| $\tau_{11\ \mathrm{sub\_ID.task}}$ | 0.59 | | |
| ICC | 0.01 | | |
| $N_{\mathrm{sub\_ID}}$ | 56 | | |
| $N_{\mathrm{stim\_ID}}$ | 539 | | |

| | |
|---|---|
| Observations | 28531 |
| Marginal $R^2$ / Conditional $R^2$ | 0.097 / 0.105 |

**TABLE S2b. Results of GLMM regression analysis predicting choice probability (online data).**

| | Choice probability for face/voice | | |
|---|---|---|---|
| Predictors | Odds Ratios | CI | p |
| (Intercept) | 0.45 | 0.38 – 0.53 | **< .001** |
| Prior cue (P- vs P=) | 0.54 | 0.39 – 0.75 | **.001** |
| Prior cue (P+ vs P=) | 1.45 | 1.05 – 2.01 | **.049** |
| Task | 0.66 | 0.49 – 0.90 | **.018** |
| Stimulus noise | 2.07 | 1.57 – 2.71 | **< .001** |
| Prior cue (P- vs P=) × task | 1.61 | 0.84 – 3.10 | .214 |
| Prior cue (P+ vs P=) × task | 1.88 | 0.98 – 3.60 | .093 |
| Prior cue (P- vs P=) × stimulus noise | 0.77 | 0.40 – 1.48 | .546 |
| Prior cue (P+ vs P=) × stimulus noise | 0.94 | 0.49 – 1.78 | .840 |
| Task × stimulus noise | 1.13 | 0.66 – 1.93 | .730 |
| Random Effects | | | |
| $\sigma^2$ | 3.29 | | |
| $\tau_{00\ stim\_ID}$ | 2.46 | | |
| $\tau_{00\ sub\_ID.2}$ | 0.56 | | |
| $\tau_{11\ sub\_ID.stimulus\_noise}$ | 0.11 | | |
| $\tau_{11\ sub\_ID.task}$ | 0.99 | | |
| ICC | 0.43 | | |
| $N_{sub\_ID}$ | 192 | | |
| $N_{stim\_ID}$ | 540 | | |
| Observations | 102739 | | |
| Marginal $R^2$ / Conditional $R^2$ | 0.059 / 0.464 | | |

### Choice behaviour as a function of stimulus category (hits and false alarms)

Simple effects of prior information on decision performance can be measured as correct and false positive target decisions (hits and false alarms [FA] respectively). Both quantities indicate how participants adapt their decisions to probabilistic cues as a function of accuracy but without accounting for bias. Both informative cues should result in the highest hit and lowest false-alarm rates across modalities.

In partial alignment with this prediction, the $P^+$ cue increased hit rates in the visual task but resulted in the lowest hit rates in the auditory task. This interaction between modalities in $P^+$ cued blocks is a result of a reduced performance benefit of this cue in the auditory task across all participants and datasets ($P^+$: $OR_{lab} = 1.73$, 95% CI = [1.47 - 2.04], $p < .001$; $OR_{online} = 2.14$ [1.96 - 2.33], $p < .001$; Tables S3; Fig. S1A). The $P^-$ cue did not improve hit rates in general in either dataset, when compared to the uninformative $P^=$ cue, which may question its use for performance ($P^-$: $OR_{lab} = 0.91$ [0.82 - 1.00], $p = .105$; $OR_{Online} = 0.91$ [0.87 - 0.96], $p = .001$; Fig. S1A). On the contrary, the $P^-$ cue appeared to reduce hit rates in the auditory, while resulting in no difference in the visual task. Our online data provided statistical evidence for the latter interaction effect of the $P^-$ cue and modality ($P^-$: $OR_{online} = 1.39$ [1.25 - 1.54], $p < .001$; Fig. S1A). Interestingly, patients' overall hit rates were significantly reduced ($OR_{main} = 0.48$ [0.27 - 0.86], $p = .030$). Hence, these results suggest a modality-specific performance benefit of informative cues on hit rates – alongside general difficulties with accurate target detection of patients with psychotic disorders.

Strategic decision adaptations towards a conservative strategy based on prior information may help to avoid false alarms (i.e., erroneously indicating a target detection when no target was present). The FA rate is significantly affected by both informative cues across participants and datasets. While we found a generally reduced FA rate in $P^-$ cued blocks ($P^-$: $OR_{lab} = 0.81$ [0.73 - 0.89], $p < .001$; $OR_{online} = 0.70$ [0.66 - 0.74], $p = .001$; Fig. S1B), the FA rate increased in $P^+$ cued blocks ($P^+$: $OR_{lab} = 1.24$ [1.11 - 1.39], $p = .001$; $OR_{online} = 1.23$ [1.16 - 1.30], $p < .001$; Fig. S1B). In the online data, both informative cues ($P^-/P^+$) also interacted significantly with modality on FA rate. That is, the $P^+$ cue resulted in more FAs in the visual compared to the auditory task ($P^+$: $OR_{online} = 1.86$ [1.65 - 2.09], $p < .001$; Fig. S1B), whereas the $P^-$ cue resulted in a reduced FA rate that is most reduced in the auditory task ($P^-$: $OR_{online} = 1.23$ [1.10 - 1.37], $p < .001$; Fig. S1B). Patients with psychosis neither exhibited a general change in FA rate nor a change specific to one cue or modality ($OR_{main} = 0.62$ [0.32 - 1.21], $p = .255$; $P^-$: $OR_{interaction} = 1.14$ [0.92 - 1.41], $p = .317$; $P^+$: $OR_{interaction} = 1.02$ [0.80 - 1.30], $p = .864$; Fig. S1B). Thus, prior information affected false positive rates across all participants, but only reached statistical significance in the well-powered online data. Together with the results on hit rates, these findings suggest cue-specific top-down modulations of decision strategy with ramifications for performance.

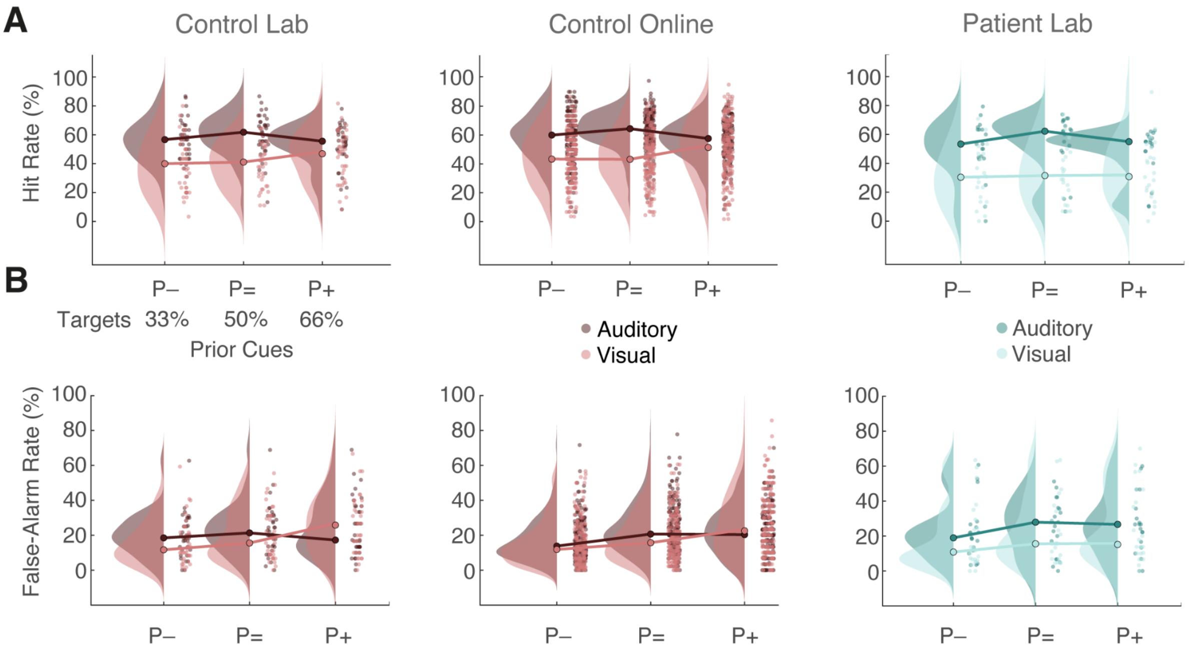


**Fig. S1. Effects of prior information on hit and false alarm rates**. Raincloud plots of **(A)** hit rate in percent and **(B)** false alarm (FA) rate in percent relative to the number of possible target (A) or non-target (B) trials, as a function of prior information cues, modality, and group. Prior information cues indicated true target probabilities, presented as $P^-$ (33%), $P^=$ (50%), and $P^+$ (66%) 'face' or 'voice' probability in the current block. Reddish colours depict data of lab and online control groups (N=36, N=192, respectively), while turquoise colours depict data of patients with psychotic disorders (N=20). Dots represent average data of individual participants.

**TABLE S3a. Results of GLMM regression analysis predicting hit rate (lab data).**

| | Hit rate | | |
|---|---|---|---|
| Predictors | Odds Ratios | CI | p |
| (Intercept) | 0.72 | 0.57 – 0.91 | **.019** |
| Chlorpromazine equivalent 100mg | 1.11 | 1.00 – 1.25 | .105 |
| Prior cue (P- vs P=) | 0.91 | 0.82 – 1.00 | .105 |
| Prior cue (P+ vs P=) | 0.98 | 0.90 – 1.07 | .704 |
| Patient | 0.48 | 0.27 – 0.86 | **.030** |
| Task | 0.45 | 0.37 – 0.54 | **< .001** |
| Stimulus noise | 0.42 | 0.37 – 0.47 | **< .001** |
| Prior cue (P- vs P=) × task | 1.18 | 0.97 – 1.44 | .156 |
| Prior cue (P+ vs P=) × task | 1.73 | 1.47 – 2.04 | **< .001** |
| Prior cue (P- vs P=) × patient | 0.95 | 0.76 – 1.18 | .704 |
| Prior cue (P+ vs P=) × patient | 0.97 | 0.81 – 1.16 | .755 |
| Prior cue (P- vs P=) × stimulus noise | 1.10 | 0.90 – 1.34 | .465 |
| Prior cue (P+ vs P=) × stimulus noise | 1.48 | 1.26 – 1.75 | **< .001** |
| Task × stimulus noise | 0.80 | 0.70 – 0.94 | **.014** |
| Patient × task | 0.72 | 0.48 – 1.07 | .156 |
| Patient × stimulus noise | 1.08 | 0.85 – 1.36 | .669 |
| Random Effects | | | |
| $\sigma^2$ | 3.29 | | |
| $\tau_{00\ sub\_ID}$ | 0.55 | | |
| $\tau_{11\ sub\_ID.task}$ | 0.40 | | |
| $\tau_{11\ sub\_ID.stimulus\_noise}$ | 0.09 | | |
| ICC | 0.14 | | |
| $N_{sub\_ID}$ | 56 | | |
| Observations | 14286 | | |
| Marginal $R^2$ / Conditional $R^2$ | 0.098 / 0.226 | | |

**TABLE S3b. Results of GLMM regression analysis predicting hit rate (online data).**

| | **Hit rate** | | |
|---|---|---|---|
| Predictors | Odds Ratios | CI | p |
| (Intercept) | 1.09 | 1.00 – 1.20 | .056 |
| Prior cue (P- vs P=) | 0.91 | 0.87 – 0.96 | **.001** |
| Prior cue (P+ vs P=) | 1.02 | 0.98 – 1.07 | .277 |
| Task | 0.53 | 0.47 – 0.60 | **< .001** |
| Stimulus noise | 2.62 | 2.48 – 2.78 | **< .001** |
| Prior cue (P- vs P=) × task | 1.39 | 1.25 – 1.54 | **< .001** |
| Prior cue (P+ vs P=) × task | 2.14 | 1.96 – 2.33 | **< .001** |
| Prior cue (P- vs P=) × stimulus noise | 0.84 | 0.76 – 0.93 | **.001** |
| Prior cue (P+ vs P=) × stimulus noise | 0.70 | 0.64 – 0.76 | **< .001** |
| Task × stimulus noise | 1.71 | 1.58 – 1.85 | **< .001** |
| Random Effects | | | |
| $\sigma^2$ | 3.29 | | |
| $\tau_{00\ sub\_ID}$ | 0.39 | | |
| $\tau_{11\ sub\_ID.task}$ | 0.65 | | |
| $\tau_{11\ sub\_ID.stimulus\_noise}$ | 0.08 | | |
| $\rho_{01}$ | 0.49 | | |
| ICC | 0.15 | | |
| $N_{sub\_ID}$ | 192 | | |
| Observations | 51371 | | |
| Marginal $R^2$ / Conditional $R^2$ | 0.086 / 0.221 | | |

**TABLE S4a. Results of GLMM regression analysis predicting false alarm rate (lab data).**

| | **False alarm (FA) rate** | | |
|---|---|---|---|
| Predictors | Odds Ratios | CI | p |
| (Intercept) | 0.19 | 0.15 – 0.25 | **< .001** |
| Chlorpromazine equivalent 100mg | 1.16 | 1.02 – 1.32 | .054 |
| Prior cue (P- vs P=) | 0.81 | 0.73 – 0.89 | **< .001** |
| Prior cue (P+ vs P=) | 1.24 | 1.11 – 1.39 | **.001** |
| Patient | 0.62 | 0.32 – 1.21 | .255 |
| Task | 0.82 | 0.65 – 1.05 | .205 |
| Stimulus noise | 0.78 | 0.71 – 0.86 | **< .001** |
| Prior cue (P- vs P=) × task | 0.93 | 0.76 – 1.13 | .593 |
| Prior cue (P+ vs P=) × task | 1.27 | 1.01 – 1.59 | .091 |
| Prior cue (P- vs P=) × patient | 1.14 | 0.92 – 1.41 | .317 |
| Prior cue (P+ vs P=) × patient | 1.02 | 0.80 – 1.30 | .864 |
| Prior cue (P- vs P=) × stimulus noise | 0.83 | 0.69 – 1.01 | .116 |
| Prior cue (P+ vs P=) × stimulus noise | 0.95 | 0.76 – 1.19 | .757 |
| Task × stimulus noise | 1.29 | 1.08 – 1.53 | **.013** |
| Patient × task | 0.89 | 0.53 – 1.50 | .757 |
| Patient × stimulus noise | 0.96 | 0.78 – 1.19 | .780 |
| Random Effects | | | |
| $\sigma^2$ | 3.29 | | |
| $\tau_{00\ sub\_ID}$ | 0.70 | | |
| $\tau_{11\ sub\_ID.task}$ | 0.67 | | |
| $\tau_{11\ sub\_ID.stimulus\_noise}$ | 0.03 | | |
| ICC | 0.17 | | |
| $N_{sub\_ID}$ | 56 | | |
| Observations | 14245 | | |
| Marginal $R^2$ / Conditional $R^2$ | 0.033 / 0.202 | | |

**TABLE S4b. Results of GLMM regression analysis predicting false alarm rate (online data).**

| | **False alarm (FA) rate** | | |
|---|---|---|---|
| Predictors | Odds Ratios | CI | p |
| (Intercept) | 0.21 | 0.19 – 0.23 | **< .001** |
| Prior cue (P- vs P=) | 0.70 | 0.66 – 0.74 | **< .001** |
| Prior cue (P+ vs P=) | 1.23 | 1.16 – 1.30 | **< .001** |
| Task | 0.86 | 0.74 – 1.01 | .086 |
| Stimulus noise | 1.15 | 1.09 – 1.23 | **< .001** |
| Prior cue (P- vs P=) × task | 1.23 | 1.10 – 1.37 | **< .001** |
| Prior cue (P+ vs P=) × task | 1.86 | 1.65 – 2.09 | **< .001** |
| Prior cue (P- vs P=) × stimulus noise | 1.12 | 1.01 – 1.25 | **.048** |
| Prior cue (P+ vs P=) × stimulus noise | 0.98 | 0.87 – 1.11 | .775 |
| Task × stimulus noise | 1.09 | 0.99 – 1.20 | .086 |
| Random Effects | | | |
| $\sigma^2$ | 3.29 | | |
| $\tau_{00\ sub\_ID}$ | 0.46 | | |
| $\tau_{11\ sub\_ID.task}$ | 1.13 | | |
| $\tau_{11\ sub\_ID.stimulus\_noise}$ | 0.06 | | |
| $\rho_{01}$ | 0.19 | | |
| ICC | 0.19 | | |
| $N_{sub\_ID}$ | 192 | | |
| Observations | 51368 | | |
| Marginal $R^2$ / Conditional $R^2$ | 0.020 / 0.203 | | |

**TABLE S5a. Results of LMM regression analysis predicting criterion (c; lab data).**

| | **Criterion** | | |
|---|---|---|---|
| Predictors | Estimates | CI | p |
| (Intercept) | 0.68 | 0.51 – 0.86 | **< .001** |
| Chlorpromazine equivalent 100mg | -0.09 | -0.17 – 0.01 | .083 |
| Task | 0.32 | 0.17 – 0.46 | **< .001** |
| Patient | 0.44 | -0.01 – 0.87 | .102 |
| Stimulus noise | -0.39 | -0.45 – -0.31 | **< .001** |
| Prior cue (P- vs P=) | 0.09 | 0.02 – 0.16 | **.024** |
| Prior cue (P+ vs P=) | -0.05 | -0.11 – 0.02 | .330 |
| Task × patient | 0.17 | -0.14 – 0.48 | .444 |
| Task × prior cue (P- vs P=) | -0.01 | -0.14 – 0.13 | .981 |
| Task × prior cue (P+ vs P=) | -0.26 | -0.39 – -0.12 | **.001** |
| Task × stimulus noise | 0.00 | -0.11 – 0.11 | .981 |
| Patient × prior cue (P- vs P=) | -0.01 | -0.15 – 0.13 | .981 |
| Patient × prior cue (P+ vs P=) | 0.02 | -0.13 – 0.16 | .981 |
| Patient × stimulus noise | 0.04 | -0.11 – 0.19 | .832 |
| Stimulus noise × prior cue (P- vs P=) | 0.00 | -0.13 – 0.14 | .981 |
| Stimulus noise × prior cue (P+ vs P=) | 0.13 | -0.01 – 0.26 | .136 |
| Random Effects | | | |
| $\sigma^2$ | 0.13 | | |
| $\tau_{00\ sub\_ID}$ | 0.29 | | |
| $\tau_{11\ sub\_ID.task}$ | 0.23 | | |
| $\tau_{11\ sub\_ID.stimulus\_noise}$ | 0.02 | | |
| ICC | 0.69 | | |
| $N_{sub\_ID}$ | 56 | | |
| Observations | 648 | | |
| Marginal $R^2$ / Conditional $R^2$ | 0.191 / 0.751 | | |

**TABLE S5b. Results of LMM regression analysis predicting criterion (c; online data).**

| | Criterion | | |
|---|---|---|---|
| Predictors | Estimates | CI | p |
| (Intercept) | 0.50 | 0.43 – 0.56 | **< .001** |
| Prior cue (P- vs P=) | -0.01 | -0.05 – 0.03 | .708 |
| Prior cue (P+ vs P=) | -0.26 | -0.30 – -0.22 | **< .001** |
| Task | 0.26 | 0.17 – 0.35 | **< .001** |
| Stimulus noise | 0.06 | 0.02 – 0.11 | **.006** |
| Prior cue (P- vs P=) × task | 0.05 | -0.04 – 0.13 | .320 |
| Random Effects | | | |
| $\sigma^2$ | 0.18 | | |
| $\tau_{00\ sub\_ID}$ | 0.17 | | |
| $\tau_{11\ sub\_ID.task}$ | 0.31 | | |
| ICC | 0.49 | | |
| N $_{sub\_ID}$ | 192 | | |
| Observations | 2304 | | |
| Marginal $R^2$ / Conditional $R^2$ | 0.076 / 0.525 | | |

**TABLE S6a. Results of LMM regression analysis predicting sensitivity (d'; lab data).**

| | Sensitivity | | |
|---|---|---|---|
| Predictors | Estimates | CI | p |
| (Intercept) | 0.87 | 0.75 – 0.98 | **< .001** |
| Chlorpromazine equivalent 100mg | -0.02 | -0.07 – 0.04 | .590 |
| Task | -0.39 | -0.55 – -0.24 | **< .001** |
| Patient | -0.22 | -0.51 – 0.06 | .186 |
| Stimulus noise | 0.42 | 0.30 – 0.53 | **< .001** |
| Prior cue (P- vs P=) | 0.13 | 0.02 – 0.24 | **.050** |
| Prior cue (P+ vs P=) | -0.07 | -0.18 – 0.04 | .280 |
| Task × patient | -0.13 | -0.46 – 0.21 | .556 |
| Task × prior cue (P- vs P=) | 0.20 | -0.02 – 0.43 | .133 |
| Task × prior cue (P+ vs P=) | 0.21 | -0.02 – 0.43 | .133 |
| Patient × prior cue (P- vs P=) | -0.16 | -0.40 – 0.07 | .254 |
| Patient × prior cue (P+ vs P=) | 0.04 | -0.20 – 0.28 | .800 |
| Patient × stimulus noise | 0.01 | -0.24 – 0.25 | .961 |
| Task × stimulus noise | 0.28 | 0.10 – 0.46 | **.010** |
| Stimulus noise × prior cue (P- vs P=) | -0.27 | -0.49 – -0.05 | **.049** |
| Stimulus noise × prior cue (P+ vs P=) | -0.37 | -0.59 – -0.14 | **.005** |
| Random Effects | | | |
| $\sigma^2$ | 0.35 | | |
| $\tau_{00\ sub\_ID}$ | 0.10 | | |
| $\tau_{11\ sub\_ID.task}$ | 0.20 | | |
| $\tau_{11\ sub\_ID.stimulus\_noise}$ | 0.06 | | |
| ICC | 0.22 | | |
| $N_{sub\_ID}$ | 56 | | |
| Observations | 648 | | |
| Marginal $R^2$ / Conditional $R^2$ | 0.220 / 0.390 | | |

**TABLE S6b. Results of LMM regression analysis predicting sensitivity (d'; online data).**

| | **Sensitivity** | | |
|---|---|---|---|
| Predictors | Estimates | CI | p |
| (Intercept) | 1.10 | 1.05 – 1.16 | **< .001** |
| Prior cue (P- vs P=) | 0.00 | -0.07 – 0.07 | .988 |
| Prior cue (P+ vs P=) | -0.09 | -0.15 – -0.02 | **.015** |
| Task | -0.25 | -0.36 – -0.14 | **< .001** |
| Stimulus noise | 0.15 | 0.08 – 0.22 | **< .001** |
| Prior cue (P- vs P=) × task | -0.18 | -0.31 – -0.04 | **.015** |
| Random Effects | | | |
| $\sigma^2$ | 0.46 | | |
| $\tau_{00\ sub\_ID}$ | 0.09 | | |
| $\tau_{11\ sub\_ID.task}$ | 0.39 | | |
| ICC | 0.17 | | |
| $N_{sub\_ID}$ | 192 | | |
| Observations | 2304 | | |
| Marginal $R^2$ / Conditional $R^2$ | 0.059 / 0.220 | | |

**TABLE S7a. Results of LMM regression analysis predicting sensitivity (d') by traits (lab data).**

| | **Sensitivity** | | |
|---|---|---|---|
| Predictors | Estimates | CI | p |
| (Intercept) | 0.70 | 0.49 – 0.91 | **< .001** |
| Chlorpromazine equivalent 100mg | -0.05 | -0.11 – 0.02 | .329 |
| Prior cue (P- vs P=) | -0.02 | -0.29 – 0.25 | .875 |
| Prior cue (P+ vs P=) | -0.14 | -0.41 – 0.12 | .425 |
| SPQ | 0.04 | 0.01 – 0.08 | .157 |
| PDI | 0.01 | -0.03 – 0.06 | .685 |
| LSHS | -0.01 | -0.03 – 0.00 | .329 |
| Patient | -0.32 | -0.80 – 0.16 | .350 |
| Task | -0.55 | -0.78 – -0.33 | **< .001** |
| Prior cue (P- vs P=) × SPQ | 0.00 | -0.04 – 0.05 | .875 |
| Prior cue (P+ vs P=) × SPQ | 0.03 | -0.02 – 0.07 | .425 |
| Prior cue (P- vs P=) × PDI | -0.03 | -0.08 – 0.02 | .382 |
| Prior cue (P+ vs P=) × PDI | -0.01 | -0.06 – 0.03 | .706 |
| Prior cue (P- vs P=) × LSHS | 0.02 | 0.00 – 0.04 | .286 |
| Prior cue (P+ vs P=) × LSHS | 0.00 | -0.02 – 0.01 | .747 |
| SPQ × patient | 0.06 | -0.02 – 0.13 | .329 |
| PDI × patient | -0.06 | -0.14 – 0.02 | .329 |
| LSHS × patient | 0.00 | -0.03 – 0.03 | .875 |
| SPQ × task | 0.03 | 0.00 – 0.07 | .286 |
| PDI × task | -0.04 | -0.08 – 0.00 | .229 |
| LSHS × task | 0.01 | -0.01 – 0.02 | .602 |
| (Prior cue (P- vs P=) × SPQ) × task | 0.02 | -0.01 – 0.05 | .382 |
| (Prior cue (P+ vs P=) × SPQ) × task | 0.03 | 0.00 – 0.06 | .286 |
| Random Effects | | | |
| $\sigma^2$ | 0.47 | | |
| $\tau_{00\ sub\_ID}$ | 0.08 | | |

| | |
|---|---|
| ICC | 0.14 |
| N $_{sub\_ID}$ | 56 |
| Observations | 648 |
| Marginal $R^2$ / Conditional $R^2$ | 0.157 / 0.273 |

**TABLE S7b. Results of LMM analysis predicting sensitivity (d') by traits (online data).**

| | Sensitivity | | |
|---|---|---|---|
| Predictors | Estimates | CI | p |
| (Intercept) | 1.15 | 1.05 – 1.26 | **< .001** |
| Prior cue (P- vs P=) | 0.03 | -0.12 – 0.18 | .829 |
| Prior cue (P+ vs P=) | -0.04 | -0.19 – 0.12 | .829 |
| SPQ | -0.01 | -0.03 – 0.01 | .740 |
| PDI | 0.00 | -0.02 – 0.03 | .920 |
| LSHS | 0.00 | -0.01 – 0.01 | .920 |
| Task | -0.28 | -0.40 – -0.15 | **< .001** |
| Prior cue (P- vs P=) × SPQ | -0.00 | -0.03 – 0.02 | .829 |
| Prior cue (P+ vs P=) × SPQ | -0.01 | -0.04 – 0.02 | .784 |
| Prior cue (P- vs P=) × PDI | -0.01 | -0.04 – 0.03 | .829 |
| Prior cue (P+ vs P=) × PDI | 0.02 | -0.02 – 0.05 | .779 |
| Prior cue (P- vs P=) × LSHS | 0.00 | -0.01 – 0.01 | .829 |
| Prior cue (P+ vs P=) × LSHS | -0.00 | -0.02 – 0.01 | .829 |
| SPQ × task | -0.03 | -0.05 – -0.01 | **.036** |
| PDI × task | 0.02 | -0.01 – 0.05 | .586 |
| LSHS × task | 0.01 | -0.00 – 0.02 | .153 |
| (Prior cue (P- vs P=) × SPQ) × task | -0.04 | -0.06 – -0.02 | **< .001** |
| (Prior cue (P+ vs P=) × SPQ) × task | -0.02 | -0.04 – -0.01 | **.041** |
| Random Effects | | | |
| $\sigma^2$ | 0.57 | | |
| $\tau_{00\ sub\_ID}$ | 0.09 | | |
| ICC | 0.13 | | |
| $N_{sub\_ID}$ | 192 | | |
| Observations | 2304 | | |
| Marginal $R^2$ / Conditional $R^2$ | 0.048 / 0.174 | | |

**TABLE S8a. Results of LMM analysis predicting criterion (c) by traits (lab data).**

| | Criterion | | |
|---|---|---|---|
| Predictors | Estimates | CI | p |
| (Intercept) | 0.79 | 0.45 – 1.12 | **< .001** |
| Chlorpromazine equivalent 100mg | -0.12 | -0.22 – 0.02 | .121 |
| Prior cue (P- vs P=) | 0.17 | -0.02 – 0.36 | .281 |
| Prior cue (P+ vs P=) | -0.07 | -0.25 – 0.12 | .805 |
| SPQ | -0.01 | -0.06 – 0.05 | .871 |
| PDI | 0.02 | -0.05 – 0.09 | .820 |
| LSHS | -0.01 | -0.04 – 0.02 | .805 |
| Patient | 0.52 | -0.24 – 1.29 | .503 |
| Task | 0.37 | 0.21 – 0.52 | **< .001** |
| Prior cue (P- vs P=) × SPQ | -0.01 | -0.04 – 0.03 | .820 |
| Prior cue (P+ vs P=) × SPQ | 0.01 | -0.03 – 0.04 | .820 |
| Prior cue (P- vs P=) × PDI | 0.00 | -0.03 – 0.04 | .910 |
| Prior cue (P+ vs P=) × PDI | 0.01 | -0.02 – 0.04 | .805 |
| Prior cue (P- vs P=) × LSHS | 0.00 | -0.02 – 0.01 | .820 |
| Prior cue (P+ vs P=) × LSHS | -0.01 | -0.02 – 0.01 | .805 |
| SPQ × patient | 0.04 | -0.09 – 0.16 | .820 |
| PDI × patient | 0.02 | -0.11 – 0.15 | .820 |
| LSHS × patient | 0.03 | -0.08 – 0.02 | .549 |
| SPQ × task | 0.02 | -0.01 – 0.05 | .399 |
| PDI × task | -0.01 | -0.03 – 0.02 | .820 |
| LSHS × task | -0.01 | -0.02 – 0.00 | .134 |
| (Prior cue (P- vs P=) × LSHS) × task | -0.01 | -0.02 – 0.01 | .741 |
| (Prior cue (P+ vs P=) × LSHS) × task | -0.01 | -0.02 – 0.00 | .053 |
| Random Effects | | | |
| $\sigma^2$ | 0.24 | | |
| $\tau_{00\ sub\_ID}$ | 0.30 | | |
| ICC | 0.56 | | |

| | |
|---|---|
| N $_{sub\_ID}$ | 56 |
| Observations | 648 |
| Marginal $R^2$ / Conditional $R^2$ | 0.142 / 0.623 |

**TABLE S8b. Results of LMM analysis predicting criterion (c) by traits (online data).**

| | Criterion | | |
|---|---|---|---|
| Predictors | Estimates | CI | p |
| (Intercept) | 0.51 | 0.38 – 0.64 | **< .001** |
| Prior cue (P- vs P=) | 0.02 | -0.09 – 0.12 | .949 |
| Prior cue (P+ vs P=) | -0.24 | -0.34 – -0.14 | **< .001** |
| SPQ | -0.01 | -0.03 – 0.01 | .841 |
| PDI | 0.00 | -0.03 – 0.03 | .956 |
| LSHS | 0.00 | -0.01 – 0.01 | .841 |
| Task | 0.25 | 0.17 – 0.34 | **< .001** |
| Prior cue (P- vs P=) × SPQ | -0.00 | -0.02 – 0.02 | .956 |
| Prior cue (P+ vs P=) × SPQ | -0.01 | -0.02 – 0.01 | .841 |
| Prior cue (P- vs P=) × PDI | -0.01 | -0.04 – 0.01 | .841 |
| Prior cue (P+ vs P=) × PDI | 0.00 | -0.02 – 0.03 | .956 |
| Prior cue (P- vs P=) × LSHS | 0.00 | -0.01 – 0.01 | .949 |
| Prior cue (P+ vs P=) × LSHS | 0.00 | -0.01 – 0.01 | .913 |
| SPQ × task | -0.02 | -0.03 – -0.00 | .065 |
| PDI × task | 0.03 | 0.01 – 0.05 | **.007** |
| LSHS × task | -0.00 | -0.01 – 0.01 | .956 |
| (Prior cue (P- vs P=) × SPQ) × task | 0.02 | -0.01 – 0.04 | .464 |
| (Prior cue (P+ vs P=) × SPQ) × task | -0.01 | -0.03 – 0.02 | .841 |
| (Prior cue (P- vs P=) × PDI) × task | -0.05 | -0.10 – -0.01 | .098 |
| (Prior cue (P+ vs P=) × PDI) × task | -0.01 | -0.06 – 0.04 | .913 |
| Random Effects | | | |
| $\sigma^2$ | 0.27 | | |
| $\tau_{00\ sub\_ID}$ | 0.17 | | |
| ICC | 0.39 | | |
| $N_{sub\_ID}$ | 192 | | |
| Observations | 2304 | | |
| Marginal $R^2$ / Conditional $R^2$ | 0.069 / 0.431 | | |

**TABLE S9. Results of LMM predicting sensitivity (d') by interviewer-rated psychosis symptoms (patient lab data).**

| | Sensitivity | | |
|---|---|---|---|
| Predictors | Estimates | CI | p |
| (Intercept) | 1.10 | 0.45 – 1.75 | **.012** |
| Chlorpromazine equivalent 100mg | -0.01 | -0.07 – 0.04 | .901 |
| PANSS positive sum | -0.01 | -0.05 – 0.02 | .844 |
| PANSS negative sum | 0.00 | -0.04 – 0.05 | .901 |
| PANSS positive sum × task | -0.03 | -0.05 – -0.02 | **< .001** |
| PANSS positive sum × prior cue (P- vs P=) | 0.00 | -0.02 – 0.03 | .901 |
| PANSS positive sum × prior cue (P+ vs P=) | 0.01 | -0.02 – 0.04 | .890 |
| PANSS negative sum × prior cue (P- vs P=) | 0.00 | -0.05 – 0.04 | .901 |
| PANSS negative sum × prior cue (P+ vs P=) | -0.02 | -0.07 – 0.02 | .835 |
| Random Effects | | | |
| $\sigma^2$ | 0.49 | | |
| $\tau_{00\ sub\_ID}$ | 0.08 | | |
| ICC | 0.14 | | |
| $N_{sub\_ID}$ | 20 | | |
| Observations | 216 | | |
| Marginal $R^2$ / Conditional $R^2$ | 0.147 / 0.266 | | |

**TABLE S10. Results of LMM regression analysis predicting criterion (c) by interviewer-rated psychosis symptoms (patient lab data).**

| | Criterion | | |
|---|---|---|---|
| Predictors | Estimates | CI | p |
| (Intercept) | 0.92 | -0.35 – 2.29 | .295 |
| Chlorpromazine equivalent 100mg | -0.09 | -0.21 – 0.02 | .295 |
| PANSS positive sum | -0.05 | -0.11 – 0.02 | .295 |
| PANSS negative sum | 0.06 | -0.03 – 0.16 | .295 |
| PANSS positive sum × task | 0.03 | 0.02 – 0.04 | **< .001** |
| PANSS positive sum × prior cue (P- vs P=) | 0.00 | -0.01 – 0.02 | .922 |
| PANSS positive sum × prior cue (P+ vs P=) | 0.00 | -0.02 – 0.02 | .922 |
| PANSS negative sum × prior cue (P- vs P=) | 0.00 | -0.03 – 0.03 | .922 |
| PANSS negative sum × prior cue (P+ vs P=) | 0.00 | -0.03 – 0.03 | .922 |
| Random Effects | | | |
| $\sigma^2$ | 0.24 | | |
| $\tau_{00\ sub\_ID}$ | 0.46 | | |
| ICC | 0.66 | | |
| $N_{sub\_ID}$ | 20 | | |
| Observations | 216 | | |
| Marginal $R^2$ / Conditional $R^2$ | 0.229 / 0.735 | | |

**TABLE S11a. Results of LMM regression analysis predicting response times (lab data).**

| | Response Time (log) | | |
|---|---|---|---|
| Predictors | Estimates | CI | p |
| (Intercept) | 7.01 | 6.93 – 7.09 | **< .001** |
| Chlorpromazine equivalent 100mg | 0.00 | -0.04 – 0.03 | .865 |
| Prior cue (P- vs P=) | -0.03 | -0.04 – -0.01 | **.001** |
| Prior cue (P+ vs P=) | -0.01 | -0.02 – 0.01 | .551 |
| Patient | 0.06 | -0.14 – 0.25 | .749 |
| Task | -0.12 | -0.17 – -0.07 | **< .001** |
| Stimulus noise | -0.01 | -0.02 – 0.01 | .666 |
| Prior cue (P- vs P=) × task | -0.01 | -0.04 – 0.01 | .551 |
| Prior cue (P+ vs P=) × task | 0.03 | 0.00 – 0.06 | .132 |
| Prior cue (P- vs P=) × stimulus noise | -0.01 | -0.03 – 0.02 | .842 |
| Prior cue (P+ vs P=) × stimulus noise | 0.02 | -0.01 – 0.04 | .551 |
| Prior cue (P- vs P=) × patient | 0.03 | 0.01 – 0.05 | **.013** |
| Prior cue (P+ vs P=) × patient | 0.01 | -0.01 – 0.03 | .667 |
| Patient × task | -0.10 | -0.20 – 0.00 | .132 |
| Patient × stimulus noise | 0.00 | -0.02 – 0.02 | .945 |
| Task × stimulus noise | 0.00 | -0.03 – 0.02 | .865 |
| (Prior cue (P- vs P=) × patient) × task | 0.04 | 0.00 – 0.08 | .132 |
| (Prior cue (P+ vs P=) × patient) × task | -0.04 | -0.08 – 0.00 | .132 |
| Random Effects | | | |
| $\sigma^2$ | 0.08 | | |
| $\tau_{00\ stim\_ID.1}$ | 0.00 | | |
| $\tau_{00\ sub\_ID.2}$ | 0.07 | | |
| $\tau_{11\ stim\_ID.patient}$ | 0.00 | | |
| $\tau_{11\ sub\_ID.stimulus\_noise}$ | 0.00 | | |
| $\tau_{11\ sub\_ID.task}$ | 0.03 | | |

| | |
|---|---|
| ICC | 0.01 |
| $N_{sub\_ID}$ | 56 |
| $N_{stim\_ID}$ | 539 |
| Observations | 28531 |
| Marginal $R^2$ / Conditional $R^2$ | 0.053 / 0.058 |

**TABLE S11b. Results of LMM regression analysis predicting response times (online data).**

| | **Response Time (log)** | | |
|---|---|---|---|
| Predictors | Estimates | CI | p |
| (Intercept) | 6.88 | 6.85 – 6.91 | **< .001** |
| Prior cue (P- vs P=) | -0.04 | -0.05 – -0.03 | **< .001** |
| Prior cue (P+ vs P=) | -0.04 | -0.05 – -0.02 | **< .001** |
| Task | -0.27 | -0.30 – -0.23 | **< .001** |
| Stimulus noise | 0.00 | -0.01 – 0.01 | .726 |
| Prior cue (P- vs P=) × task | 0.00 | -0.03 – 0.03 | .906 |
| Prior cue (P+ vs P=) × task | 0.05 | 0.02 – 0.08 | **.001** |
| Prior cue (P- vs P=) × stimulus noise | 0.02 | -0.01 – 0.05 | .230 |
| Prior cue (P+ vs P=) × stimulus noise | -0.02 | -0.05 – 0.01 | .230 |
| Task × stimulus noise | -0.02 | -0.04 – 0.01 | .230 |
| Random Effects | | | |
| $\sigma^2$ | 0.07 | | |
| $\tau_{00\ stim\_ID}$ | 0.00 | | |
| $\tau_{00\ sub\_ID.2}$ | 0.03 | | |
| $\tau_{11\ sub\_ID.stimulus\_noise}$ | 0.00 | | |
| $\tau_{11\ sub\_ID.task}$ | 0.05 | | |
| ICC | 0.06 | | |
| $N_{sub\_ID}$ | 192 | | |
| $N_{stim\_ID}$ | 540 | | |
| Observations | 102739 | | |
| Marginal $R^2$ / Conditional $R^2$ | 0.201 / 0.247 | | |

**TABLE S12. Results starting point (*z*).** Bold indicates an overlap of less or more than 20% of a distribution with the mean of the comparator distribution. Distributions depicted in Fig. 5C-D.

| **Group** | **Modality** | **Cue (prior information)** | **Mean** | **2.5q** | **97.5q** | **P(cue < \| > 0.5)** | **P($P^+$ > M $P^-$)** |
|---|---|---|---|---|---|---|---|
| Control | A | P+ | 0.547 | 0.525 | 0.569 | **1** | 0.352 |
| Control | A | P= | 0.510 | 0.486 | 0.533 | NA | NA |
| Control | A | P– | 0.512 | 0.490 | 0.536 | **0.163** | NA |
| Control | V | P+ | 0.538 | 0.516 | 0.563 | **1** | **0.937** |
| Control | V | P= | 0.518 | 0.495 | 0.543 | NA | NA |
| Control | V | P– | 0.491 | 0.468 | 0.514 | 0.78 | NA |
| Patient | A | P+ | 0.542 | 0.502 | 0.578 | **0.983** | 0.421 |
| Patient | A | P= | 0.511 | 0.474 | 0.551 | NA | NA |
| Patient | A | P– | 0.496 | 0.458 | 0.532 | 0.566 | NA |
| Patient | V | P+ | 0.489 | 0.454 | 0.524 | 0.269 | 0.660 |
| Patient | V | P= | 0.481 | 0.445 | 0.515 | NA | NA |
| Patient | V | P– | 0.467 | 0.432 | 0.504 | **0.965** | NA |

**TABLE S13. Results drift criterion/evidence accumulation bias (dc).** Bold indicates an overlap of less or more than 20% of a distribution with the mean of the comparator distribution. Distributions depicted in Fig. 5E-H.

| Group | Modality | Cue (prior information) | Mean | 2.5q | 97.5q | P(cue > M $P^{=}$) | P(min overlap distributions) |
|---|---|---|---|---|---|---|---|
| Control | A | P+ | -0.471 | -0.639 | -0.303 | **0.954** | **0.046 ($P^{=}$)** |
| Control | A | P= | -0.27 | -0.435 | -0.102 | NA | NA |
| Control | A | P– | -0.427 | -0.593 | -0.263 | **0.907** | NA |
| Control | V | P+ | -0.614 | -0.779 | -0.450 | **0.118** | **0.063 ($P^{-}$)** |
| Control | V | P= | -0.756 | -0.924 | -0.591 | NA | NA |
| Control | V | P– | -0.805 | -0.975 | -0.636 | 0.661 | NA |
| Patient | A | P+ | -0.454 | -0.747 | -0.161 | 0.276 | 0.276 ($P^{=}$) |
| Patient | A | P= | -0.333 | -0.626 | -0.041 | NA | NA |
| Patient | A | P– | -0.408 | -0.712 | -0.122 | 0.368 | NA |
| Patient | V | P+ | -0.689 | -0.962 | -0.421 | 0.557 | 0.340 ($P^{-}$) |
| Patient | V | P= | -0.713 | -0.983 | -0.440 | NA | NA |
| Patient | V | P– | -0.77 | -1.048 | -0.495 | 0.385 | NA |

**TABLE S14a. HDDM model comparisons DIC – participants without psychosis.**

| Model | Parameters | DIC |
|---|---|---|
| M0c | full model | 31000 |
| M1c | fixed starting point | 31212 |
| M2c | fixed drift criterion | 31311 |

**Table S14b. HDDM model comparisons DIC – patients with psychosis.**

| Model | Parameters | DIC |
|---|---|---|
| M0p | full model | 18278 |
| M1p | fixed starting point | 18311 |
| M2p | fixed drift criterion | 18436 |
